# Light-driven repair: Photobiomodulation restores blood–brain barrier function following hypoxic injury

**DOI:** 10.64898/2026.02.15.706027

**Authors:** Mirriam Domocos, Denis E. Bragin, Nagesh Shanbhag, Luise Schlotterose, Mootaz M. Salman

## Abstract

A functional blood-brain barrier (BBB) is essential for the central nervous system (CNS) homeostasis and its disruption is an early event in acute brain injury and chronic neurodegeneration. Hypoxia triggers BBB breakdown, promoting endothelial dysfunction, oxidative stress, metabolic dysregulation and thrombo-inflammatory responses that compromise barrier integrity. However, strategies that directly restore BBB function remain limited. Here, we investigated whether photobiomodulation (PBM), a non-invasive light therapy, can rescue BBB dysfunction following acute hypoxic stress. Using a multicellular in vitro BBB model comprising immortalised human brain microvascular endothelial cells, pericytes and astrocytes, we induced hypoxic injury (6 h, 1% O₂) and applied three PBM treatments during recovery. Hypoxia significantly reduced transendothelial electrical resistance (TEER), whereas PBM restored barrier function in endothelial monocultures and tri-cultures. Endothelial cells showed the strongest hypoxic response, with increased hypoxia-inducible factor-1α, plasminogen activator inhibitor-1 and von Willebrand factor (vWF), all attenuated by PBM. Importantly, siRNA-mediated knockdown of vWF partially recapitulated PBM-induced barrier rescue, identifying endothelial vWF as a mediator of recovery. PBM also reduced reactive oxygen species in hypoxic astrocytes and pericytes, indicating coordinated multicellular modulation. These findings demonstrate that PBM restores BBB integrity after hypoxic insult by modulating endothelial thrombo-inflammatory signalling while reducing oxidative stress in glial cells. Rather than acting as a general cytoprotective stimulus, PBM engages defined molecular pathways linked to endothelial activation. This work establishes a mechanistically informed platform for studying BBB repair and supports PBM as a targeted strategy to protect vascular integrity in hypoxia-associated neurological disorders.

**Key points summary:**

- Hypoxia is a major driver of blood-brain barrier (BBB) dysfunction, yet there are currently no targeted therapies that directly restore barrier integrity.
- Photobiomodulation (PBM) is a non-invasive low-level light intervention known to facilitate mitochondrial function and cellular stress responses.
- In a human in vitro BBB model, repeated PBM treatment restored transendothelial electrical resistance (TEER) 24 and 48 hours after hypoxic injury, with endothelial rescue linked to downregulation of von Willebrand factor (vWF).
- PBM modulated oxidative stress, hypoxia signalling, and thrombo-inflammatory pathways across endothelial cells, astrocytes, and pericytes.
- These findings support light-driven modulation of endothelial signalling as a potential strategy to restore BBB integrity in hypoxia-associated neurological conditions.

**Graphical abstract:** 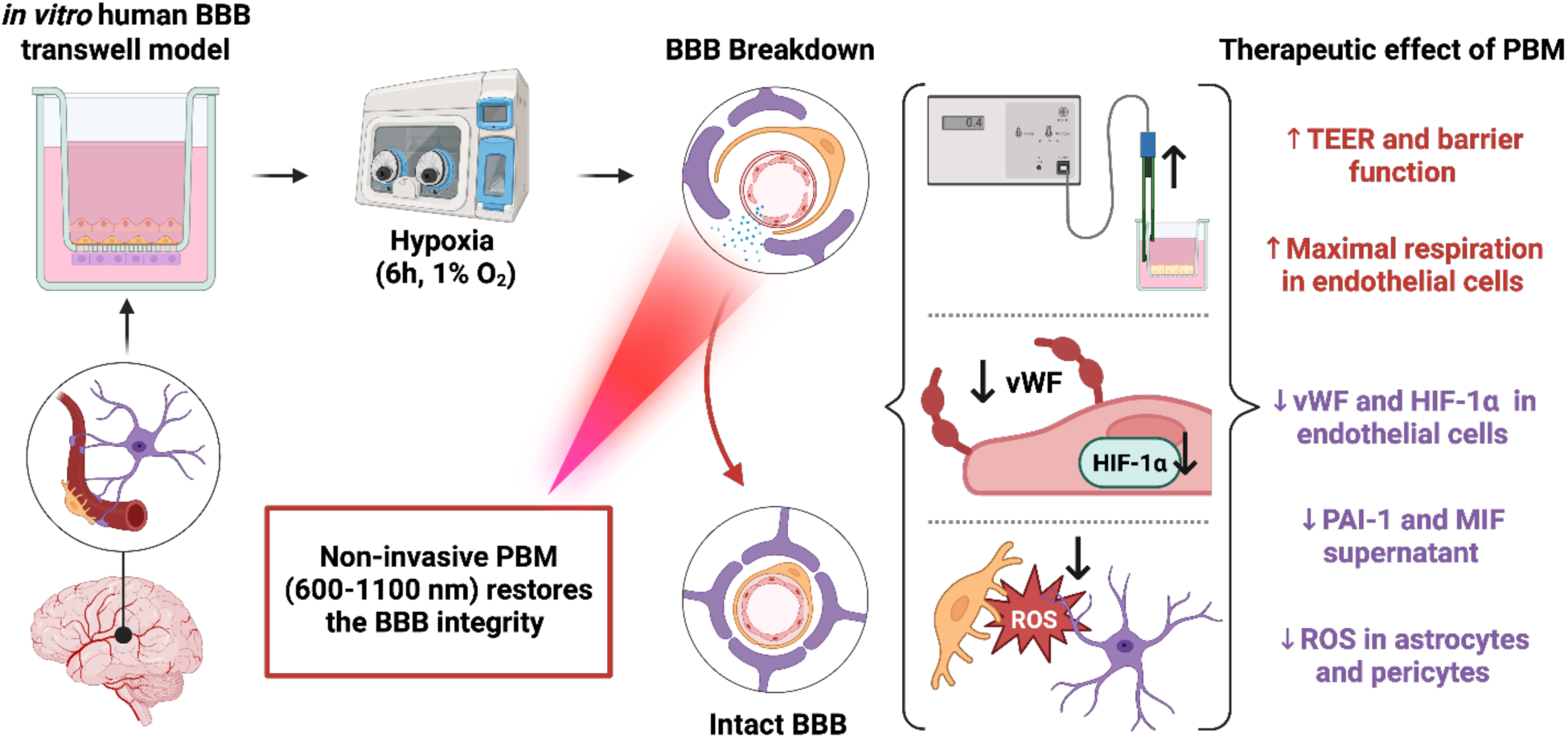

## 1. Introduction

The blood-brain barrier (BBB) is a highly specialised and dynamically regulated multicellular interface that preserved central nervous system (CNS) integrity by controlling nutrient and oxygen supply, coordinating metabolic waste clearance, and preventing the entry of potentially neurotoxic circulating factors [1]. The brain endothelial cells make up a continuous monolayer lining the blood vessel, distinguished from peripheral endothelial cells by high expression of tight junction proteins, most importantly zonula occludens-1 (ZO-1), claudin-5, and occludin [2, 3]. Wrapped around endothelial cells are pericytes that give structural support as well as maintain the BBB by the expression of the platelet-derived growth factor receptor beta (PDGFRβ). Brain pericytes on larger vessels express higher levels of alpha smooth muscle actin (αSMA) to regulate the blood flow and oxygenation of the CNS [3, 4]. The astrocytes complete the BBB by “grabbing” the vessel with their flattened endfeet extension which allows for close communication between the vasculature and neurons. An essential protein of the endfeet structure is the water channel aquaporin-4 (AQP4) that facilitates bidirectional water movement through the brain [5, 6]. Given the importance of the BBB in brain homeostasis, damage to the BBB is associated with a variety of disease, including dementias and stroke, and leads to the structural breakdown of tight junctions and loss of endothelial cells, pericytes, and astrocytes. This disrupts normal nutrient transport, increases BBB permeability leading to a leaky barrier, and triggers inflammatory responses that cause neuronal damage, impaired waste clearance, and brain and cell death [7, 8]. Hypoxia is a common source of BBB dysfunction where low oxygenation stabilises hypoxia-inducible factor-1 alpha (HIF-1α), which translocates to the cell nucleus and facilitates inflammatory responses, metabolic reprogramming, and vascular remodelling. In addition, hypoxia leads to oxidative stress, characterised by mitochondrial dysfunction and upregulation of reactive oxygen species (ROS) [9–12]. In endothelial cells, the platelet adhesion molecule von Willebrand factor (vWF) becomes upregulated and contributes to the infiltration of leukocytes and a decrease in tight junction proteins [13]. Pericytes and astrocytes detach from the basal membrane, leaving the BBB unsupported and thus downregulate key cell markers linked to BBB function. Overall, hypoxic- and HIF-1α-mediated mechanisms impair the barrier integrity and with a lack of therapeutic interventions this leaves a vicious cycle of BBB dysfunction and disease progression [14].

Photobiomodulation (PBM) is a non-invasive and FDA (Food and Drug administration)-approved low-level light therapy (LLLT) that exhibits neuroprotective effects and boosts the cell’s metabolism and adenosine triphosphate (ATP) synthesis while reducing inflammation and oxidative stress. Commonly used in dermatology, sports injury, pain, and tissue healing, the red and infrared light ranges from 600 to 1100 nm and becomes absorbed by the fourth terminal enzyme in the mitochondria, the cytochrome C oxidase (CCO) [15]. PBM additionally modulates oxygen (O_2_) free radicals and ROS levels [16].

Further studies have reported beneficial effects of PBM on cognitive defects and anxiety in APP/PS1 transgenic in vivo models, including an upregulation of ZO-1, claudin-5, and occludin which improved the BBB integrity [17]. The clinical significance of PBM has been previously shown on endothelial dysfunction, suggesting improved recovery after coronary intervention and maintenance of endothelial function by harmonising endothelial nitric oxide (eNO) and transforming growth factor beta (TGF-β) levels [18]. Despite growing interest in PBM as a neuroprotective strategy, its cell-specific effects on the BBB remain poorly defined, and it is unclear whether PBM can directly restore barrier integrity following hypoxic injury.

To address this gap, this present study aimed to determine the therapeutic impact of PBM on BBB function under hypoxic condition and to dissect the intrinsic responses of individual cellular components of the BBB. We employed a controlled in vitro BBB model consisting of immortalised microvascular human brain endothelial cells co-cultured with human astrocytes and vascular pericytes in a transwell system. This reduction is yet physiologically informed platform enabled precise interrogation of barrier function and facilitated downstream mechanistic assays to define cell-type–specific responses.

Our findings demonstrate that three sessions of five-minute PBM irradiation significantly restored the barrier function of endothelial monocultures and BBB tri-cultures following hypoxic insult (6 hours at 1%). This functional recovery was associated with a downregulation of hypoxia-induced vWF in endothelial cells, attenuation of HIF-1α expression, and enhancement of maximal mitochondrial respiratory capacity indicating improved metabolic resilience. Alongside, PBM reduced ROS levels in hypoxic astrocytes and pericytes, highlighting coordinated multi-cellular effects. Cytokine and chemokine profiling further revealed modulation of thrombo-inflammatory pathways consistent with a protective anti-thrombotic milieu. Collectively, these results identify previously unrecognised mechanisms which PBM non-invasively re-establishes BBB function after hypoxic injury and support its potential as a therapeutic strategy for cerebrovascular dysfunction.

## 2. Materials and Methods

### 2.1. Cell culture lines

The cell lines used were purchased from Innoprot: human brain microvascular endothelial cells (HBMECs) (Innoprot; P10361-IM), human astrocytes (HAs) (Innoprot; P10251-IM), and human brain vascular pericytes (HVPCs) (Innoprot; P10363-IM). The cell lines were immortalised via the SV40 antigen T. Cells were used up to passage number 20 and maintained at 37°C and 5% CO_2_ in the appropriate medium purchased from Innoprot, with a full medium change every two days: endothelial cell medium (EM) (Innoprot; P60104), astrocyte medium (AM) (Innoprot; P60101), or pericyte medium-PLUS (PM) (Innoprot; P60121-Plus).

### 2.2. Culture for transwell models

Cell-to-cell contact models were established to facilitates communication between cells on Millicell® 24-well hanging inserts (Millipore; PTSP24H48) with a pore size of 3 µm and area of 0.3 cm^2^. On day 1, the basal sides of the inserts were inverted, resting inside the cover plate lid, and coated with 50 µL of Poly-L-lysine (PLL) (2 µg/cm^2^) (Sigma-Aldrich; P4705). A cover plate lid was placed on top to distribute the coating, and the plate was incubated at room temperature (RT) for 1 h. PLL coatings were washed with PBS (Gibco; 10010023) and for models with HAs, 20 µL of HA suspension (1.5 × 10^4^ cells) was evenly pipetted on the basal site of the membrane. The bottom of the well plate was inverted without damaging any inserts, and the cells were incubated at 37°C for 2 hours. The entire plate was flipped back with the basal side facing down, and each well was topped up with 1.3 mL AM.

On day 2, the apical sides of the inserts were coated with 200 µL of Geltrex^TM^ (Thermo Scientific; A1413302) and incubated at 37°C for 1 hour. Geltrex was then fully aspirated and for cultures with HVPCs, 100 µL of HVPC suspension (2.5 × 10^4^ cells) was pipetted into the insert, incubated for 10 minutes. For models with HVPCs and HBMECs co-cultured at the apical site, 100 µL of HBMEC suspension containing 7.5 × 10^4^ cells was used, for a total of 1.0 × 10^5^ cells seeded within the insert. 500 µL of EM:PM media was added as 1:1. For models with only HBMECs on the apical site, 100 µL of HBMEC suspension with 1.0 × 10^5^ cells was pipetted and 500 µL of EM added. The cultures were incubated at 37°C until day 8 when conditioning took place.

### 2.3. Experimental goups

On day 8, transwell models were either kept as an untreated control group 1) normoxia, or exposed to 2) PBM treatment, 3) hypoxia (6 hours at 1% O_2_), or 4) hypoxia followed by PBM. The topwrap TW-03B (ProNeuroLight, LLC, Phoenix, AZ, USA) was used to irradiate the cultures with red and infrared light in 3 sessions either, under normoxia or after hypoxia. The plate was positioned 10 cm above the flat-laid topwrap, and the transwell models were irradiated for 5 minutes. Between the sessions, plates were incubated at 37°C for a recovery period of 1 hour. PBM treatments were performed with the main light turned off, and the device placed away from the window. An EDTA-treated (50 mM) (Invitrogen; 15575-020) tri-culture acted as a negative control of barrier breakdown for all four groups. Functional and quantitative analyses were carried out 24 and 48 hours after conditioning on day 9 or 10 (Figure 1A).

**Figure 1.**
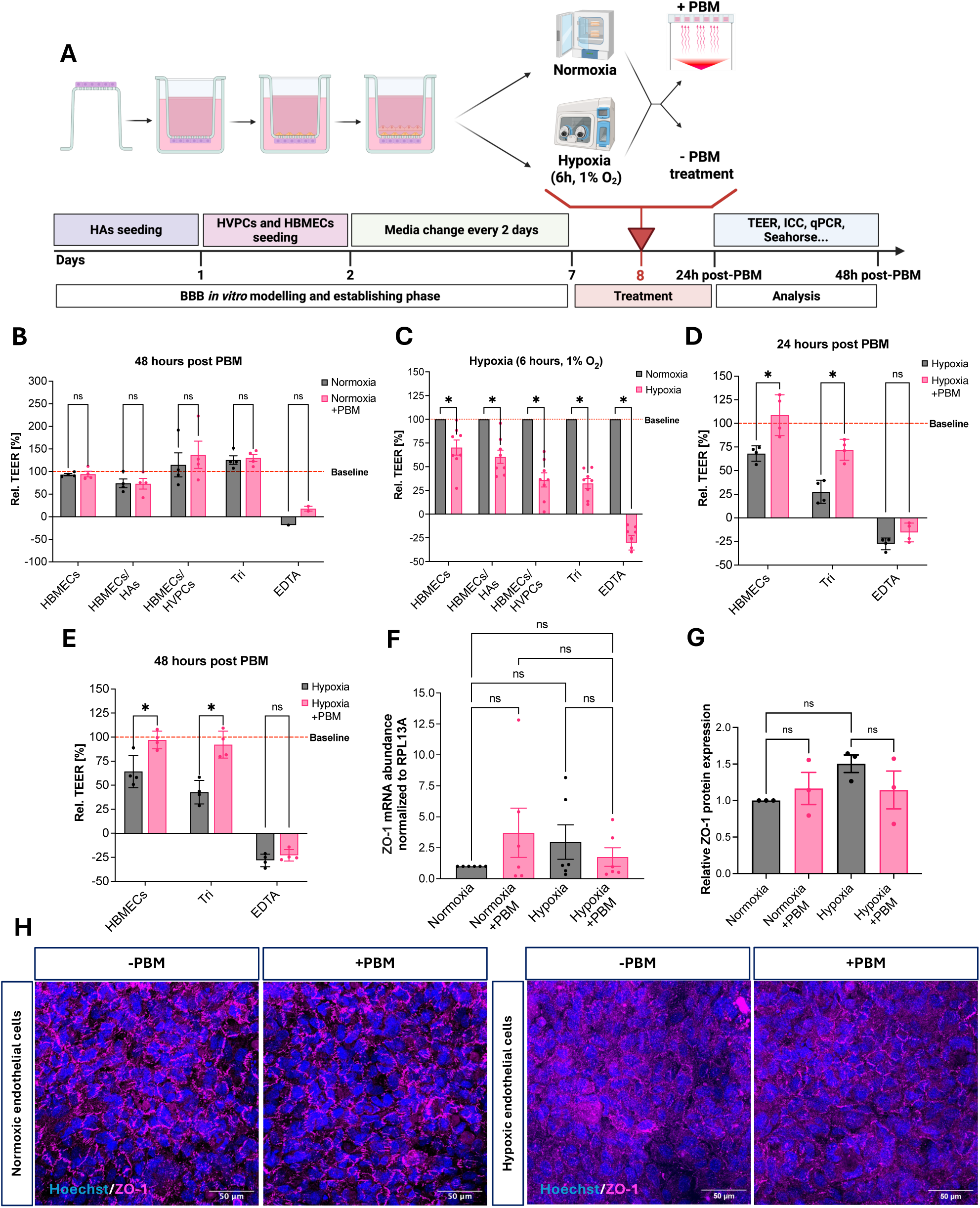
PBM restores endothelial junctional integrity and barrier function following hypoxic injury. A) Experimental workflow illustrating the transwell-based BBB model, hypoxic exposure (6 h, 1% O₂), PBM treatment schedule, and downstream functional and molecular analyses (created with BioRender). (B–E) TEER expressed as relative change (%) from baseline. (B) Normoxic controls at 48 h. (C)TEER immediately following hypoxia. (D)TEER at 24 h post-hypoxia. (E)TEER at 48 h post-hypoxia. PBM significantly restored endothelial barrier resistance under hypoxic conditions. Statistical comparisons were performed using multiple unpaired t-tests with Holm–Šídák correction; N=4 independent biological replicates; *p<0.05. (F)ZO-1 mRNA expression in HBMECs at 48 h, normalised to RPL13A. (G)Quantification of ZO-1 protein levels relative to control, measured as ZO-1-positive area normalised to Hoechst nuclear area. (H)Representative ICC images showing ZO-1 (magenta) and nuclei (Hoechst, blue) in normoxic and hypoxic HBMECs with and without PBM. For mRNA and protein analyses, statistical significance was determined using one-way ANOVA with Šídák’s post hoc correction; N=3–4 independent biological replicates. Data are presented as mean±SEM.

### 2.4. Transendothelial electrical resistance (TEER) measurements

TEER was calculated based on the resistance measured with the EVOM^TM^ Manual Meter (World Precision Instruments; EVM-MT-03-02), following the manufacturer’s instruction for handling and measuring. Electrodes were positioned using the electrode placement frame, with the shorter electrode in the insert and the longer outside the transwell. Per transwell, three individual resistance measurements were recorded after being referenced to the blank. The TEER is the product of the membrane’s area (0.3cm^2^) and the referenced resistance. TEER was then averaged and taken relative to the baseline TEER to calculate the change in relative TEER. Measurements were taken in four biological replicates.

### 2.5. ToxiLight® cytotoxicity assay

The ToxiLight Non-Destructive Cytotoxicity BioAssay Kit (Lonza; LT17-217) was used following the manufacturers protocol and done for the hypoxia groups after 48 hours. Cell supernatant from the insert (20 µL) was transferred to a white 96-well plate with clear bottoms with two technical replicates per sample. 100 µL of the reconstituted adenylate kinase detection reagent was added to each well and incubated for 5 minutes. The luminescence was measured with the CLARIOstar PLUS microplate reader and the relative average luminescence (RLU) calculated by dividing the luminescence of samples by that of their respecive cell-free media. To calculate the relative luminescence the endpoint RLU was divided by the baseline RLU. Measurements were taken in four biological replicates with every two technical replicates.

### 2.6. AlamarBlue^TM^ cell viability assay

The alamarBlue^TM^ reagent assay (Invitrogen; DAL1025) was used to detect cell viability in normoxia and hypoxia groups at baseline (day 8) and after 48 hours. 10% of the apical and basal volumes of the alamarBlue dye were added to the appropriate compartments and incubated at 37°C for 2 hours. The sample was pipetted into a black 96-well microplate at 10% of the apical and basal volume. The fluorescence was measured at excitation/emission of 495/519 nm with the CLARIOstar PLUS microplate reader and the relative fluorescence signal was calculated by taking the average of the fluorescence from both compartments, after subtracting the background signal from the blank (the media), relative to the baseline fluorescence signal before any conditioning. Measurements were taken in four biological replicates.

### 2.7. Immunocytochemistry (ICC)

Transwell cultures were fixed with 4% Paraformaldehyde (PFA) (Thermo Scientific; 28908) in both compartments and washed twice with PBS for 5 minutes. Cells were permeabilised with 0.3% Triton X-100 (Invitrogen; HFH10) for 10 minutes at RT and washed twice again. Blocking was done with 0.5% BSA (Sigma-Aldrich; A8806-1G) and 0.5% Glycine (Sigma-Aldrich; G8898-500G) for 1 hour at RT. Primary antibodies (Table 1) were diluted appropriately in blocking solution (BS), and cells were incubated in both compartments at 4°C overnight. After washing twice for 10 minutes, cells were incubated with secondary antibodies (Table 1) and Hoechst (Invitrogen; H2569) (0.8 µL per 1 mL) diluted in BS for 1.5 hours. Cells were washed twice for 5 minutes, and membranes were cut out with a scalpel and mounted with Fluoromount (Millipore; 345789). Cells were imaged with the 3i Marianas Confocal Microscope (Intelligent Imaging Innovations) at 63X and analysed via Fiji Image J 1.54p with Java 21.0.7 (64-bit). Analysis was performed on three biological replicates, with three representative images per replicate

**Table 1:** Primary and secondary antibodies used for staining of endothelial cells, astrocytes, and pericytes.

| Antibody | Host | Dilution | Catalog number | Supplier |
| --- | --- | --- | --- | --- |
| ZO-1 | Rat | 1:200 | 15652 | Cell Signaling |
| vWF | Sheep | 1:400 | ab11713 | Abcam |
| AQP4 | Rabbit | 1:200 | ab128906 | Abcam |
| GFAP | Rat | 1:500 | 13-0300 | Invitrogen |
| PDGFRb | Rabbit | 1:200 | MA5-15143 | Invitrogen |
| HIF-1α | Mouse | 1:200 | MA1-516 | Thermo Scientific |
| Actinred™ 555 | - | 2 drops/mL | R37112 | Life Technologies Ltd |
| Anti-Sheep 647 | Donkey | 1:1000 | A21448; LOT: 2720396 | Invitrogen |
| Anti-Rat 488 | Donkey | 1:1000 | A21208; LOT: 2668657 | Invitrogen |
| Anti-Mouse 555 | Donkey | 1:1000 | A31570; LOT: 3034154 | Invitrogen |
| Anti-Rabbit 647 | Donkey | 1:1000 | A31573; LOT: 2997084 | Invitrogen |
| Anti-Mouse 488 | Donkey | 1:1000 | A2120; LOT: 3006753 | Invitrogen |

### 2.8. RNA extraction and quantitative real-time PCR analysis

HBMECs, HVPCs, and HAs were plated in 12-well plates (3.0 × 10^5^ cells), cultured to a confluency of 80%, and then conditioned for each group. After 48 hours, total RNA was extracted with TRIzol (Zymo; R2050-1-200) and purified with the Direct-zol RNA MicroPrep kit (ZYMNO Research; R2062). RNA was quantified with the NanoDrop One/One^C^ spectrophotometer (Thermo Scientific). To generate 20 µL cDNA (500 ng), 1 µL Random Primers (1:20) (Invitrogen; 48190011) and 1 µL 10 mM dNTP mix (Invitrogen; 18427089) were added to each diluted RNA sample for a final volume of 13 µL. The mixture was heated at 65 °C for 5 minutes, then cooled to 4°C for at least 1 minute. The reverse transcription was accomplished using 1 µL SuperScript III RT (200 U/µL), 4 µL 5X First Strand Buffer, 1 µL 0.1 M DTT, included in the SuperScript III RT kit (Invitrogen; 18080093), and 1 µL RnaseOUT Recombinant Ribonuclease Inhibitor (40 U/µL) (Invitrogen; 10777019) under the following conditions: 65°C for 5 minutes, 55°C for 60 minutes, and 70°C for 15 minutes. One quantitative polymerase chain reaction (qPCR) reaction (20 µL) contained 10 µL 1x Fast SYBR Green Master Mix (Applied Biosystems; 4385612), 1 µL of each the relevant forward and reverse primers (0.2 µM) (Table 2), 5 µL of cDNA template (1:10), and 3 µL dH_2_O. *RPL13A* was used as the housekeeping (HK) gene. Reactions were run with the QuantStudio 3 Real-time PCR System (Applied Biosystems; A28567) and the qPCR data was analysed following the 2^−ΔΔCT^ method normalising to the untreated normoxic cells as the control. The assay was measured in four to six biological replicates.

**Table 2:**
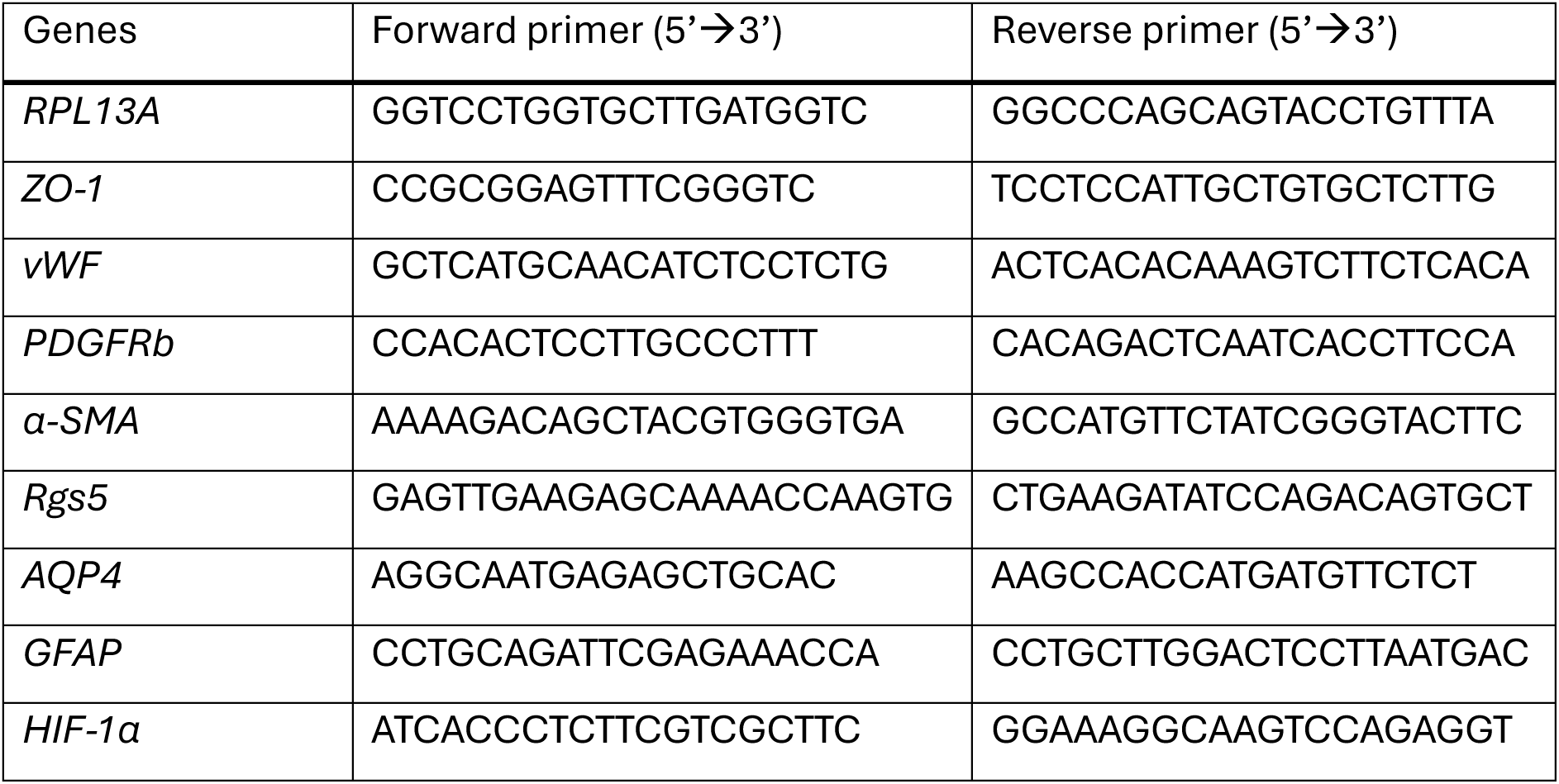
Forward and reverse sequences for all primers used in 5’ to 3’.

### 2.9. siRNA transfection and knockdown

To knock down vWF, HBMECs (5×10^5^ cells) were plated in Geltrex-coated 6-well plates and cultured for two days. Pre-annealed and pre-designed siRNA (Life Technologies Ltd; AM16708) was used to target vWF, according to the manufacturers protocol. For one well, 9 µL of Lipofectamine^®^ RNAiMAX Reagent (Life Technologies Ltd; 13778-075) was mixed with 150 µL of Opti-MEM^®^ Medium (Gibco; 31985062). 3 µL of reconstituted siRNA (Working solution: 10 µM) was added to 150 µL of Opti-MEM^®^ Medium. The dilutions were then mixed 1:1 and incubated at RT for 5 minutes. 250 µL of the siRNA-lipid complex was added to HBMECs, and the cells were incubated at 37°C for 48 hours. To investigate the pathway involvement of vWF, four biological replicates of siRNA-treated HBMECs were cultured. To confirm the knockdown of vWF, 5×10^5^ treated HBMECs were centrifuged, and the pellet was subjected to further qPCR analysis.

### 2.10. ROS-Glo^TM^ H_2_O_2_ assay

The ROS-Glo^TM^ H_2_O_2_ assay (Promega; G8820) was used to analyse the hydrogen peroxide (H_2_O_2_) production for normoxia and hypoxia groups at baseline, after one, two, and three irradiations of PBM, and after 48 hours following the manufacturer’s protocol. Cells were plated in 96-well plates (1.5 × 10^5^ cells), cultured to a confluency of 80%, and then conditioned for each group. 20 µL of the prepared H_2_O_2_ substrate solution was added and the cells were incubated at 37°C for 1 hour. 100 µL of the supernatant was transferred to a white 96-well plate, and 100 µL of the prepared ROS-Glo^TM^ detection solution was added. After incubating at RT for 20 minutes the luminescence was measured with the CLARIOstar PLUS microplate reader. Measurements were taken in four biological replicates with three technical replicates.

### 2.11. Seahorse real-time cell metabolic analysis

HBMECs, HVPCs, and HAs were plated on a Seahorse XF^e^ 96 cell culture plate (3.0 × 10^5^ cells) (Agilent; 103794-100) and the assay was carried out for normoxia and hypoxia groups after 24 and 48 hours, following the manufacturer’s protocol. Cells were cultured for two days before being conditioned for each group. On the day of the assay, cells were washed twice with 50 µL of the prepared assay media (40 mL of XF base medium (Agilent; 103575-100), 0.0728 g D glucose (10 mM) (Sigma-Aldrich; G7021), 0.00448 g sodium pyruvate (1 mM) (Sigma-Aldrich; P5280-25G), and 400 µL L-glutamine (2 mM) (Life Technologies; 25030024)). Assay media (175 µL) was added and cells were incubated at 37°C in a CO_2_-free incubator for 1 hour. After calibration of the Seahorse XF^e^ 96 Extracellular Flux Analyzer (Agilent), firstly three baseline recordings were made, followed by five after 1 µM Oligomycin (Sigma-Aldrich; O4876-5MG) treatment, three after 1 µM FCCP (Carbonyl cyanide-4 (trifluoromethoxy) phenylhydrazone) (Sigma-Aldrich; C2920-10MG) injection, three after 0.5 µM Rotenone/Antimycin A (R/A) (Sigma-Aldrich: R8875-1G;A867a) injection, and five after 50 mM 2-Deoxy-Glucose (2-DG) (Sigma-Aldrich; D8375-10MG) treatment. The oxygen consumption rate (OCR) and extracellular acidification rate (ECAR) were normalised to each well’s total protein count quantified via the Pierce^TM^ BCA protein assay kit (Thermo Scientific; 23225). Measurements were taken in four biological replicates with ten technical replicates.

### 2.12. Proteome profiler human cytokine array

The relative expression levels of 36 cytokines, chemokines, and acute phase proteins associated with inflammation were detected using the Proteome Profiler Human Cytokine Array Kit (R&D Systems; ARY005B), under normoxia and hypoxia groups after 48 hours following the manufacturers protocol. Membranes were blocked at RT for 1 hour and incubated at 4°C overnight with the sample mixture containing 1 mL cell-free supernatant taken from tri-cultures of each group, 500 µL of assay buffer 4, and 15 µL of human cytokine array detection antibody cocktail. After washing three times, 2 mL of Streptavidin-HRP (1:2000) was added and incubated at 25°C for 30 minutes on a shaker. Chemi reagents 1 and 2 were mixed 1:1 and 1 mL pipetted on washed membranes before incubating at RT for 1 minute. The membranes were imaged with the iBright Imager 1500 (Invitrogen) auto-exposure system, and the mean intensity for each dot was quantified with Fiji Image J 1.54p. Measurements were taken in one biological replicate with two technical replicates.

### 2.13. Statistics

Statistical analyses were performed using GraphPad Prism (version 10.0). Data are presented as mean ± standard error of the mean (SEM), based on a minimum of three independent biological replicates per condition. Comparisons between two independent groups were analysed using a two-tailed unpaired Student’s t-test (with equal variance assumed unless variance testing indicated otherwise). For comparisons involving more than two groups, one-way ANOVA followed by Tukey’s post hoc test was used. Where multiple pairwise comparisons across experimental conditions were required, multiple t-tests were conducted with Holm–Šídák correction to control for family-wise error. Statistical significance was defined as *p < 0.05, **p < 0.01, ***p < 0.001, and ****p < 0.0001.

## 3. Results

### 3.1. PBM restores hypoxia-induced barrier dysfunction in optimised in vitro BBB models

TEER measurements provide a direct, quantitative readout of barrier function and are widely used to assess the integrity and tightness of the BBB. Figure 1A is a schematic representation of the workflow starting at day one and two with cell seeding followed by the appropriate treatment or conditioning on day eight. To optimise the transwell model, PLL, Geltrex, and Matrigel coatings on the apical side were tested. TEER measurements revealed that Geltrex facilitates the highest barrier function in HBMEC monocultures (**p<0.01) and co-cultured with HAs (*p<0.05) (Figure S1A). PBM irradiation did not affect the TEER for normoxia groups at 48 hours (Figure 1B), thus cultures were exposed to hypoxia to explore PBM’s potential therapeutic effect in disease. Hypoxia conditions (6 hours at 1% O_2_) were selected based on prior literature [19, 20] and showed a significant decrease in TEER of around 30-70% under hypoxia (*p<0.05) across all cultures compared to normoxic cultures (Figure 1C). In HBMEC monocultures and BBB tri-cultures PBM led to a significant recovery close to baseline levels in TEER after hypoxia (*p<0.05) after 24 (108.7%±10.82 and 72.15%±5.524) and 48 hours (97.06%±4.594 and 92.21%±7.029) compared to non PBM-treated cultures (68.10%±4.048 and 27.63%±6.057 at 24 hours; 64.33%±8.411 and 42.71%±6.108 at 48 hours) (Figures 1C, D). Next, to conclude whether the restoration of barrier function was driven by the upregulation of tight junction proteins or other molecular mechanisms, the gene and protein expression of ZO-1 in HBMEC monocultures were analysed. RNA expression data showed no statistical difference between non-PBM and PBM-treated normoxia and hypoxia conditions (Figure 1F). Upon ICC, ZO-1 distribution and localisation were clearly disrupted after hypoxia compared to normoxia groups (Figure 1H). While the quantitative analysis also showed no significant difference (Figure 1G), PBM appeared to reorganise ZO-1 and improve localisation after hypoxia compared to untreated hypoxia HBMECs. Altogether, these findings reveal the therapeutic effect of PBM on HBMEC monocultures and BBB tri-cultures following hypoxic injury.

### 3.2. PBM modulates hypoxia-induced oxidative stress and HIF-1α signalling in a cell-type–specific manner

Given the mechanism of action of PBM via the CCO and thus modulation of O_2_, the synthesis of ROS from the mitochondria and interrelated HIF-1α levels after hypoxia were examined. ROS levels were measured at a “baseline level” before any treatment and after one, two, and three rounds of light irradiation, as well as after 48 hours for each cell type. After hypoxia, HAs immediately responded to one PBM treatment with a significant halving in ROS levels, from around 1752 RLU to 857.2 RLU, which remained consistent throughout all following irradiations (**p<0.01) (Figure 2B). Likewise, hypoxic HVPCs manifested with a statistical decrease of around 50% in ROS production after three rounds of PBM (*p<0.05) (Figure 2C). While ROS levels were not lowered in hypoxic HBMECs upon PBM (Figure 2A), HIF-1α was downregulated, as seen in ICC imaged in Figure 2D. To quantify HIF-1α levels from ICC, the mean intensity of HIF-1α co-localised with the cell nuclei was taken, as hypoxia allows the translocation to the nucleus for downstream pathways. In hypoxic HBMECs, the quantitative analysis revealed a significant attenuation (**p<0.01) of almost three quarters of HIF-1α after 48 hours (Figure 2D). Although not statistically relevant, it seemed that PBM also decreased HIF-1α in hypoxic HAs (p=0.0692) (Figure 2E). PBM did not alter HIF-1α levels in HVPCs (Figure 2F). Coherent with the TEER measurements, PBM did not impact the levels of ROS and HIF-1α in normoxia (Figure S2) but modulated them cell-specifically after hypoxia which could be one of the mechanisms leading to an improved BBB function.

**Figure 2.**
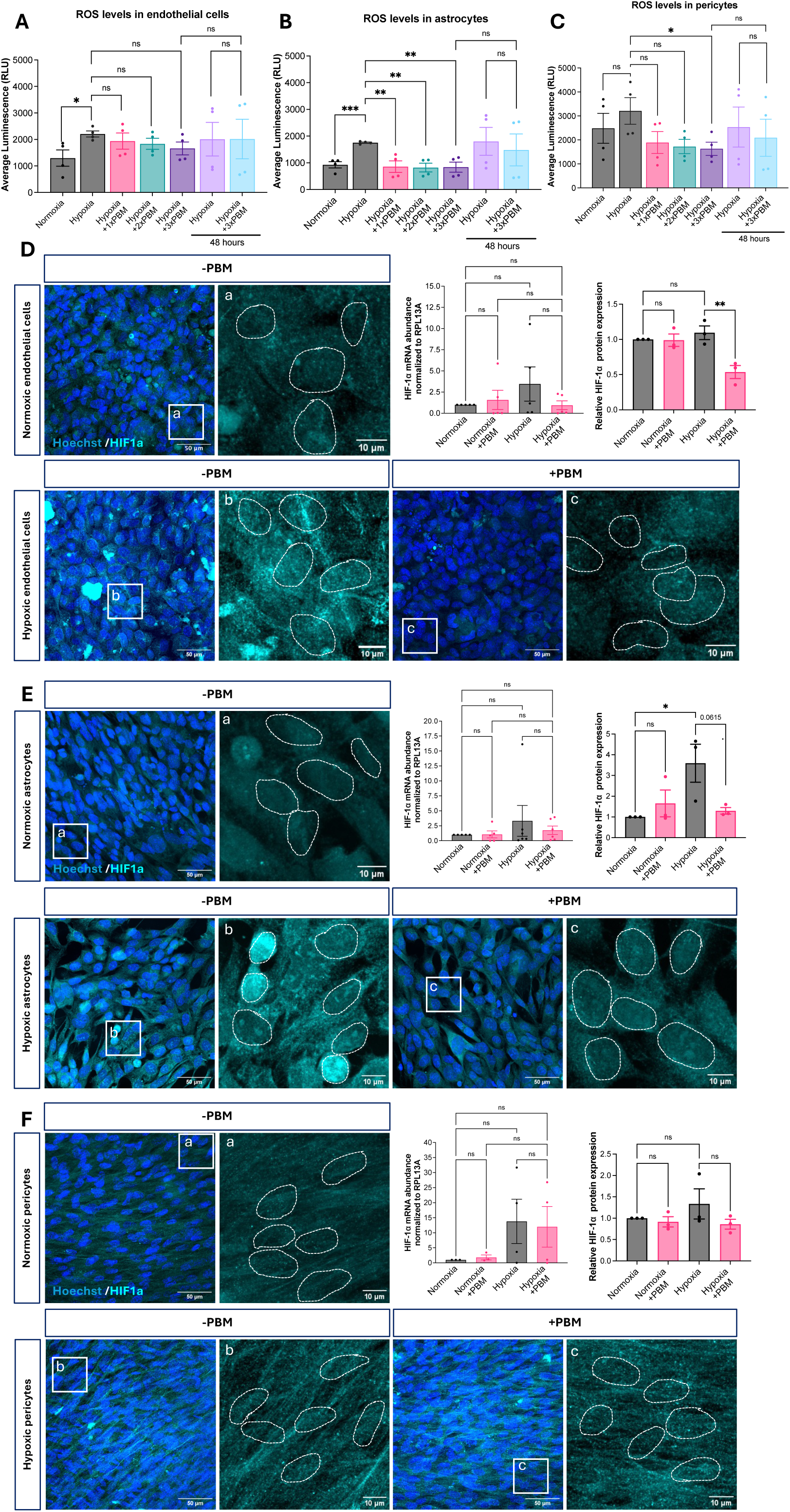
PBM attenuates oxidative stress and suppresses HIF-1α stabilisation following hypoxic exposure. (A-C) Quantification of ROS levels measured by luminescence assay in endothelial cells (A), astrocytes (B), and pericytes (C) following hypoxia (6 h, 1% O₂) with or without PBM treatment. PBM significantly reduced hypoxia-induced ROS accumulation in astrocytes and pericytes, with a modest effect in endothelial cells. (D-F) Representative ICC images showing HIF-1α (cyan) and nuclei (Hoechst, blue) in endothelial cells (D), astrocytes (E), and pericytes (F) under normoxia (–PBM), hypoxia (–PBM), and hypoxia with PBM (+PBM). Insets display higher-magnification views illustrating nuclear localisation of HIF-1α. The mRNA expression of HIF-1α was normalised to RPL13A. Nuclear HIF-1α protein levels were quantified as mean nuclear fluorescence intensity normalised to Hoechst area and expressed relative to normoxic controls. Statistical comparisons were performed using one-way ANOVA with Šídák’s post hoc correction; N=3–4 independent biological replicates. Data are presented as mean±SEM.

### 3.3. PBM increases mitochondrial respiratory capacity in endothelial cells

ROS and suppressed HIF-1α signalling, we next examined whether these effects were accompanied by changes in mitochondrial bioenergetic function in endothelial cells, astrocytes, and pericytes. Cells in normoxia and hypoxia groups were treated with oligomycin, FCCP, R/A, and 2-DG after 24 and 48 hours and the OCR and ECAR were measured with the Seahorse XF^®^ 96 Extracellular Flux Analyzer. Figure 3 visualizes OCR over 24 hours after hypoxia, showing a clear, inconsistent, and aberrant pattern of O_2_ consumption compared to the normoxia groups, which slightly normalised by 48 hours (Figure S4A-C). The basal respiration was significantly reduced by 58.46% in HBMECs (***p<0.001) and 69.29% in HAs (*p<0.05) at 24 hours after hypoxia compared to the normoxic cells (Figure 3A, B). At 48 hours after hypoxia, cells did fully recover to the control baseline (Figure S4A, B). The ATP-linked respiration was calculated as the difference between the basal respiration and the oligomycin injection and was strongly impaired 24 hours after hypoxia. Yet, there appeared to be a trend toward PBM treatment almost restoring ATP-linked respiration in HBMECs, compared to the untreated hypoxic group. This seemed to be a pattern among the HAs and HVPCs as well. Hypoxia significantly impacted the maximal respiration in HBMECs (*p<0.05) and HVPCs (*p<0.05). PBM irradiation significantly boosted the drop in maximal respiration of HBMECs after hypoxia nearly by four times compared to non-PBM hypoxic HBMECs (*p<0.05), calculated as the measurements after FCCP injection. The proton leak and spare capacity in hypoxic cells after 24 hours did not change with PBM (Figure S3), and neither did any of the OCR-linked respiration after 48 hours, as cells appeared to recover from hypoxia (Figure S4D-F).

**Figure 3.**
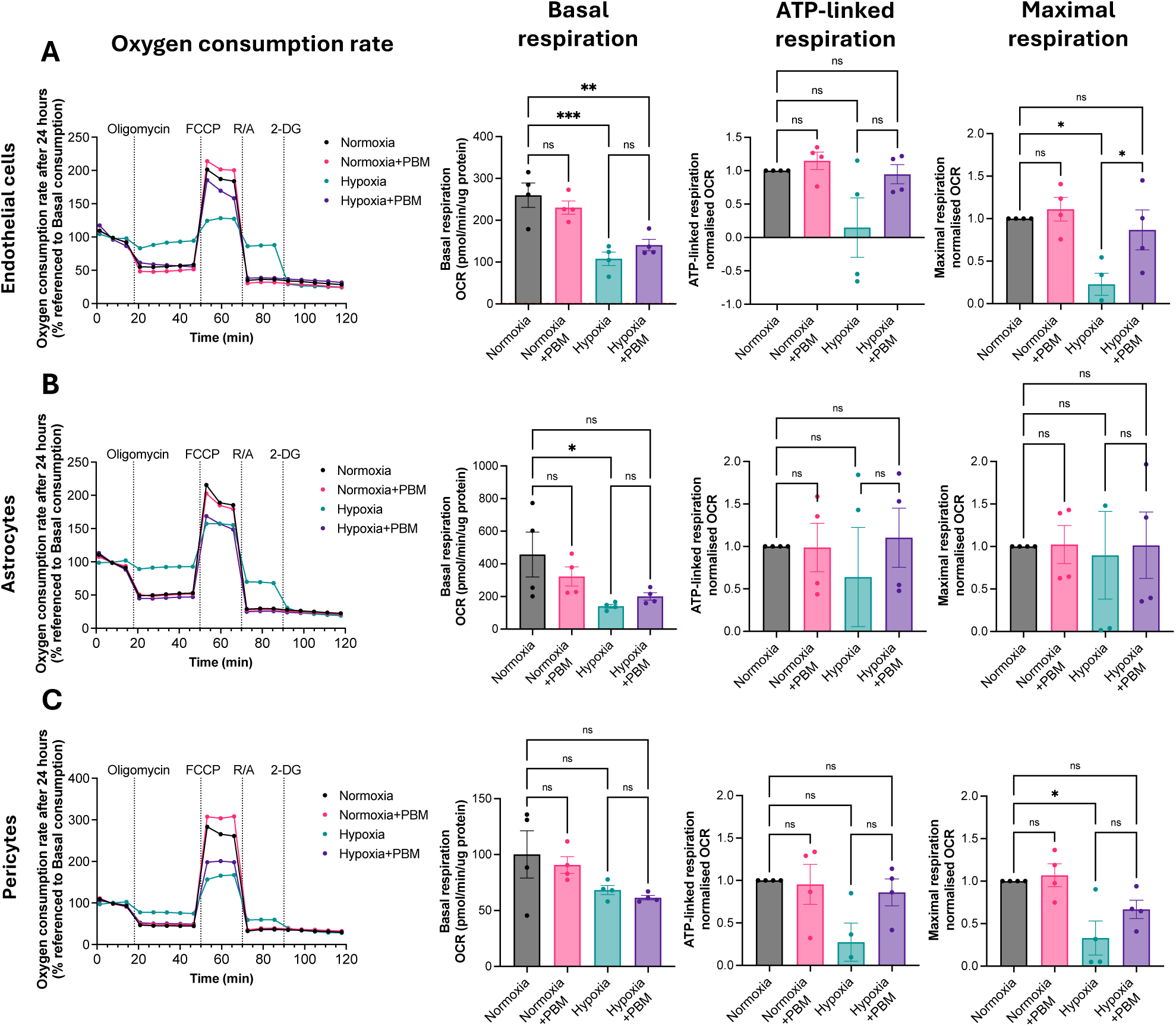
Hypoxia impacts mitochondrial respiration in endothelial cells and astrocytes, while PBM enhances maximal respiratory capacity in endothelial cells. (A–C) Mitochondrial oxygen consumption rate (OCR) profiles measured 24 hours after hypoxia (6 h, 1% O₂) in in endothelial cells (A), astrocytes (B), and pericytes (C), with or without PBM treatment. OCR traces are plotted as percentage change over time, normalised to protein content and baseline respiration. Sequential injections of oligomycin and FCCP were used to determine bioenergetic parameters. Basal respiration was calculated prior to oligomycin injection (pmol/min/µg protein). ATP-linked respiration was determined as the reduction in OCR following oligomycin and expressed relative to baseline OCR. Maximal respiration was calculated following FCCP treatment and normalised to OCR. Hypoxia significantly reduced mitochondrial function in endothelial cells and astrocytes, whereas PBM selectively increased maximal respiratory capacity in endothelial cells under hypoxic conditions. Statistical comparisons were performed using one-way ANOVA with Šídák’s post hoc correction; N=4 independent biological replicates. Data are presented as mean±SEM.

Figure S5 covers the ECAR for normoxia and hypoxia groups at 24 and 48 hours. Overall, hypoxia did not change the basal glycolytic rate after 24 hours compared to normoxia (Figure S5A-C). Interestingly, after 48 hours glycolysis decreased significantly below the normoxic control for HBMECs (**p<0.01) and HVPCs (*p<0.05) following hypoxia. The reserved glycolytic capacity was calculated as the difference between the basal glycolytic rate and the oligomycin injection. At both timepoints PBM did not alter the glycolytic rate or reserved glycolytic capacity within normoxia and hypoxia groups. Conclusively, hypoxia impacted the oxygen-linked respiration of HBMECs and HAs after 24 hours which recovered after 48 hours. Interestingly, although overall metabolic activity returned toward baseline levels, the glycolytic rate was reduced. In hypoxic endothelial cells, PBM significantly increased maximal respiratory capacity at 24 hours, indicating enhanced mitochondrial reserve and suggesting improved capacity for ATP production under stress conditions.

### 3.4. PBM alters cytokine and chemokine networks in hypoxic BBB model

PBM exhibits anti-inflammatory effects and since cells acutely respond to hypoxia by activating inflammatory pathways, it was next investigated whether PBM modulates cytokines, chemokines, and acute phase proteins in the BBB. The relative protein expression of 36 human inflammatory markers was analysed in the supernatant of tri-cultures in all four conditions at 48 hours via the Proteome Profiler Human Cytokine Array (Figure 4A). Six out of the 17 detected markers (Table S1) revealed a significant difference between the untreated and PBM-treated groups (Figure 4B). Under normoxic conditions, PBM significantly upregulated chemokines interleukin-8 (IL-8) (***p<0.001; Figure 4C) and growth-regulated oncogene-α (CXCL1/GROα) (***p<0.001; Figure 4F) as well as cytokines interleukin-21 (IL-21) (***p<0.001; Figure 4D) and granulocyte-macrophage colony-stimulating factor (GM-CSF) (***p<0.001; Figure 4G) compared to the untreated group. Following hypoxia, PBM also led to the upregulation of IL-8 (*p<0.05) and IL-21(*p<0.01), but not of CXCL1/GROα and GM-CSF. The plasminogen activator inhibitor-1 (Serpin E1/PAI-1) (Figure 4E), a pro-thrombotic protein, and migration inhibitory factor (MIF) (Figure 4H), a pro-inflammatory cytokine, were both significantly downregulated upon PBM irradiation in normoxia (***p<0.001) and hypoxia (*p<0.05) compared to non-PBM groups. Collectively, these data reveal that PBM impacts the immune regulation and response independently of BBB impairment. These changes could have protective effects, such as reduced blood clot formation and promoted angiogenesis, that lead to the repair of the damaged BBB after oxidative stress.

**Figure 4.**
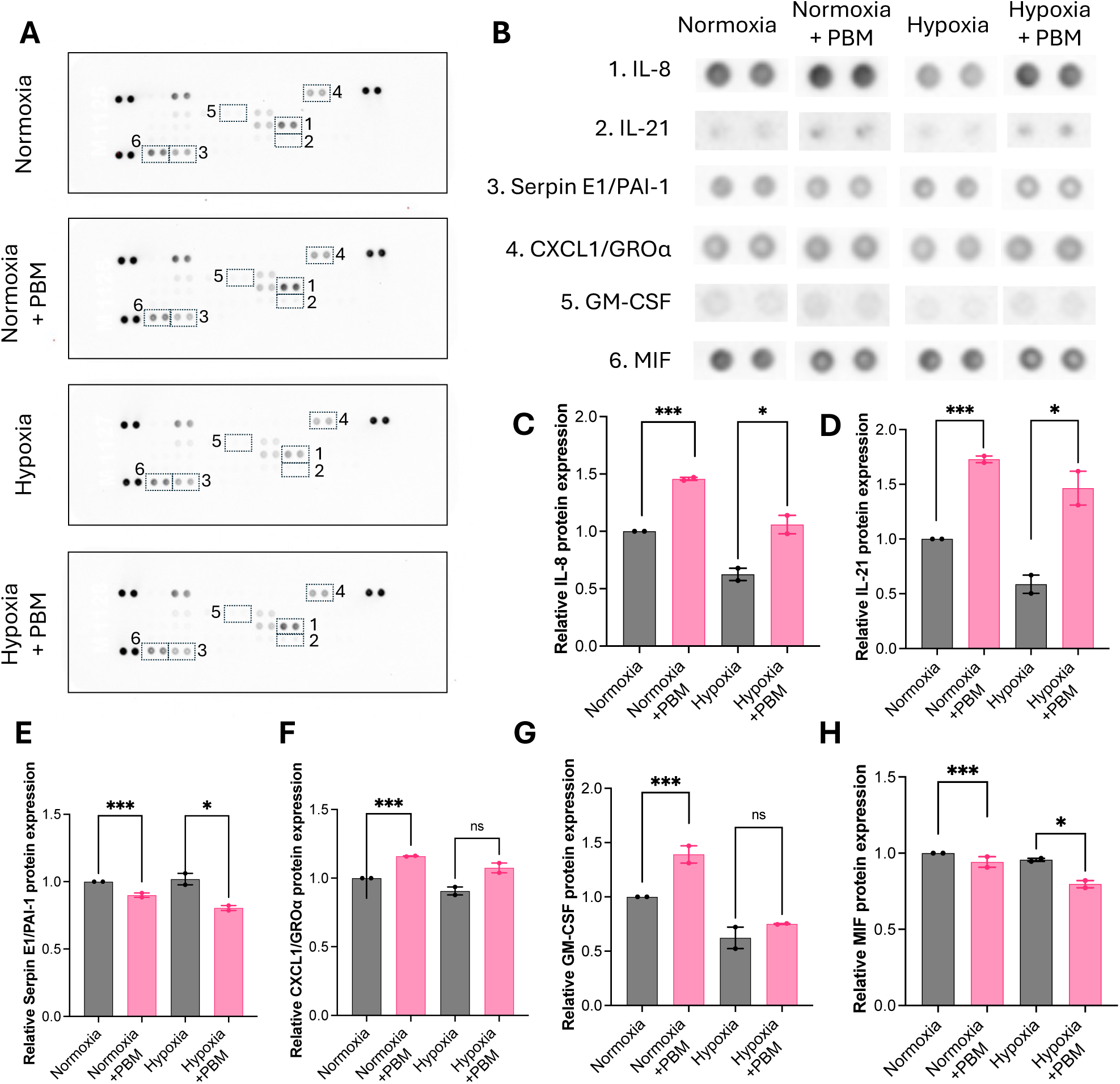
PBM differentially modulates cytokine and chemokine secretion in BBB tri-cultures following hypoxia. (A)Representative cytokine array membranes from BBB tri-culture supernatants under normoxia and hypoxia (6 h, 1% O₂), with or without PBM treatment. Seventeen markers were detected above background. (B)Quantitative summary of the six cytokines/chemokines significantly altered by PBM treatment. Spot intensities were background-subtracted and normalised to the normoxia −PBM condition. (C-H) Relative protein levels of IL-8 (C), IL-21 (D), Serpin E1/PAI-1 (E), CXCL1/GROα (F), GM-CSF (G), and MIF (H). PBM significantly downregulated the pro-thrombotic factor PAI-1 and the pro-inflammatory cytokine MIF under both normoxic and hypoxic conditions, while selectively modulating IL-8, IL-21, CXCL1, and GM-CSF. Statistical analysis was performed using unpaired two-tailed t-tests; with at least two technical replicates. Data are presented as mean±SEM.

### 3.5. vWF modulation disrupts barrier integrity and abrogates the restorative effect of PBM on TEER in endothelial cells

The cytokine array revealed an association between upregulated IL-8 levels and HBMECs. Weibel-Palade bodies (WPBs), the same granules that store vWF, also serve as a reservoir for IL-8. Given the involvement of vWF during hypoxia, its expression patterns in HBMECs were assessed. RNA expression of vWF revealed a drastic upregulation in non PBM-treated HBMECs at 48 hours after hypoxia (****p<0.0001) compared to normoxia groups (Figure 5A). The same effect was also observed upon ICC after 48 hours as seen in Figure 5B. Most strikingly, PBM significantly alleviated this effect by 85.63% in hypoxic HBMECs, nearly returning it to control levels (****p<0.0001), as detected by qPCR (Figure 5A). Quantitative analysis of the mean intensity of vWF in ICC validated these findings on a protein level with PBM-treated hypoxic HBMECs having reduced vWF expression compared to untreated hypoxic cells (*p<0.05) (Figure 5C). To evaluate whether vWF’s downregulation is involved in improving the barrier function, HBMECs were transfected with siRNA against vWF with Lipofectamine^TM^ RNAiMAX. The almost complete knock down of vWF was confirmed via qPCR (****p<0.0001) (Figure 5D). Transfected HBMECs were next used to seed monocultures and tri-cultures to assess the TEER for all four conditions at 24 and 48 hours. Cultures with depleted vWF, showed no significant change in TEER between non-PBM and PBM-treated cultures in normoxia (Figure 5E) and hypoxia (Figure 5F) at both time points. Thereby, it can be concluded that the PBM-triggered downregulation of vWF in hypoxic HBMECs is directly linked to the improvement of barrier integrity, manifesting with an increase in TEER.

**Figure 5.**
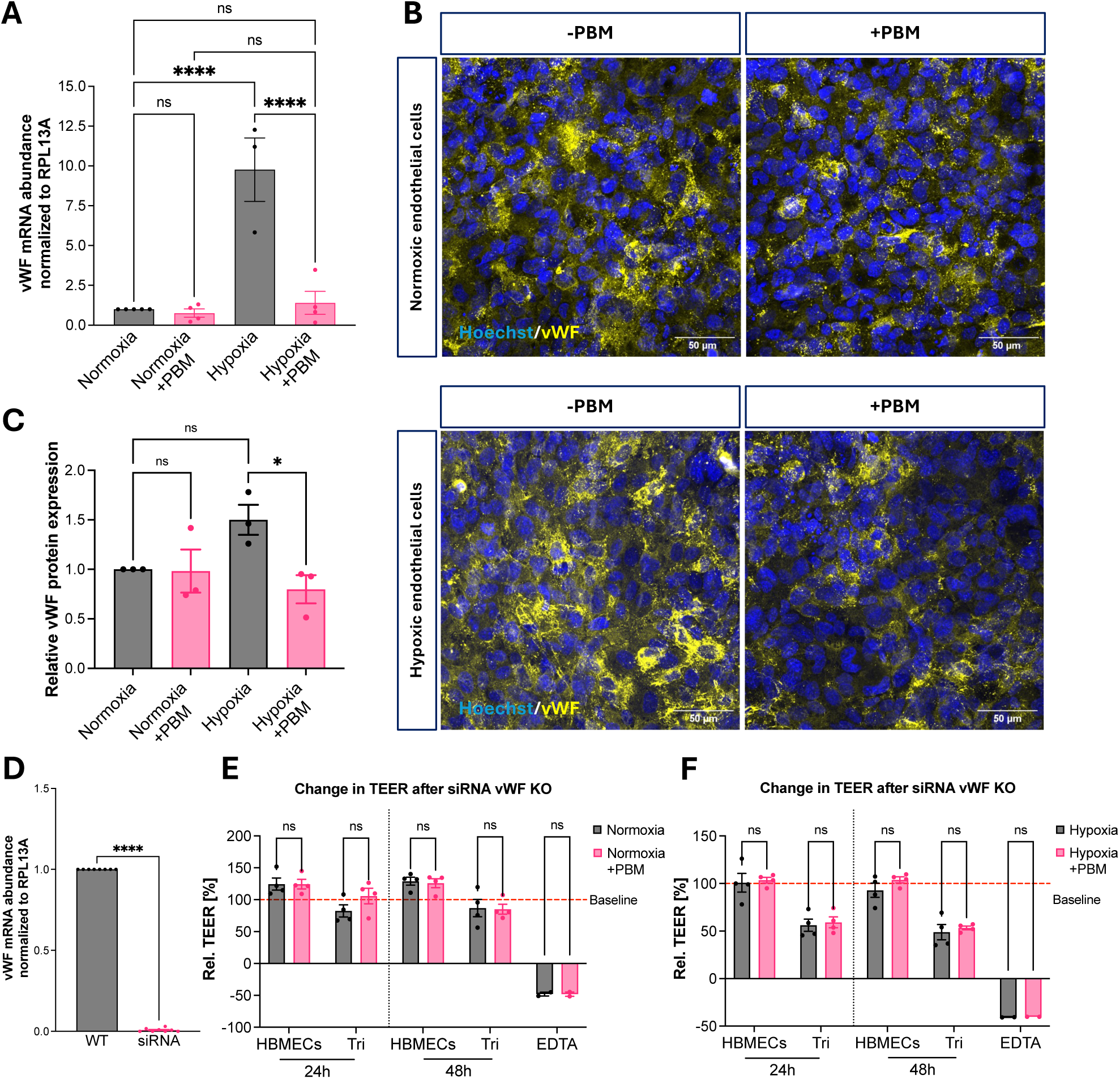
PBM restores barrier integrity through downregulation of endothelial vWF. (A)vWF mRNA expression in HBMECs 48 h after normoxia or hypoxia (6 h, 1% O₂), with or without PBM treatment, normalised to RPL13A. (B)Representative *ICC* images showing vWF (yellow) and nuclei (Hoechst, blue) in normoxic and hypoxic endothelial cells ±PBM. (C)Quantification of vWF protein levels, expressed as mean nuclear-normalised fluorescence intensity relative to normoxia −PBM controls. (D)Validation of vWF knockdown efficiency in siRNA-transfected endothelial cells, shown as relative vWF mRNA expression normalised to RPL13A. (E-F) Relative TEER changes (%) in endothelial monocultures and BBB tri-cultures at 24 h and 48 h under normoxic (E) and hypoxic (F) conditions following vWF silencing. Statistical comparisons for mRNA and protein expression were performed using one-way ANOVA with Šidák’s post hoc test (N=3–4 biological replicates). siRNA validation was analysed using an unpaired two-tailed t-test (N=8 biological replicates). TEER data were analysed using multiple unpaired t-tests with Holm–Šidák correction (N=4 biological replicates). Data are presented as mean±SEM.

### 3.6. PBM does not primarily act through classical astrocyte/pericyte reactivity markers

To delineate the cell-specific contributions of perivascular astrocytes and pericytes to endothelial barrier integrity, we next examined how these cells respond to hypoxia and whether PBM modulates these responses. Astrocytes are known to undergo reactive changes under hypoxic stress, including activation of neuroinflammatory pathways marked by glial fibrillary acidic protein (GFAP) upregulation, cellular swelling, and alterations in aquaporin-4 (AQP4) localisation. Nevertheless, the qPCR data (Figure S6A) and quantitative analysis of ICC (Figure S6C) between normoxic and hypoxic HAs revealed no significant upregulation, even though there appeared to be a trend in upregulation in the ICC images (Figure S6B). PBM treatment did not alleviate this trend in hypoxic HAs compared to non-PBM hypoxic cells. Similarly, the quantitative analysis of hypoxic HAs stained for AQP4 (Figure S6F) revealed no significant difference in the mean intensity compared to the control group. Moreover, PBM did not statistically affect expression at the RNA (Figure S6D) or protein (Figure S6F) levels under normoxia or hypoxia. Yet the intensity of AQP4 in ICC images of hypoxic HAs seemed to be less enhanced than in non-PBM hypoxic cells. Overall, these data indicate that PBM does not restore BBB integrity through modulation of classical astrocytic reactivity markers, as neither AQP4 nor GFAP expression was significantly altered under the conditions tested.

To date, there is limited evidence addressing how human vascular pericytes respond to PBM under normoxic or hypoxic conditions. To address this gap, we examined the expression of PDGFRβ, a canonical pericyte marker critically involved in pericyte survival, endothelial–pericyte signalling, and maintenance of BBB integrity. Using ICC (Figure S7B), hypoxia clearly diminished PDGFRβ levels and altered its localisation in HVPCs compared to normoxic groups. This observation was supported by measuring the mean intensity at a 50% reduction in PDGFRβ, although the difference was not statistically significant (Figure S7C). Within the normoxia and hypoxia groups PBM did not alter the expression and localisation of PDGFRβ. Since the RNA expression was below the detectable range, the expression of Rgs5, its upstream regulator during hypoxia, was quantified in Figure S7A. In contrast to the literature, Rgs5 was found to be downregulated in hypoxic HVPCs, suggesting that the reduction in PDGFRβ may depend on alternative pathways under hypoxia.

Finally, to assess functional alterations in HVPCs across conditions, we examined cytoskeletal organisation and αSMA dynamics under hypoxia and PBM treatment. Pericyte contractility, largely governed by actin filaments and αSMA, plays a key role in regulating microvascular tone and supporting endothelial barrier stability. Although hypoxia did not significantly alter αSMA mRNA expression compared to normoxia, and PBM had no detectable transcriptional effect within groups (Figure S7D), structural changes were evident at the cytoskeletal level. Immunostaining revealed that hypoxia disrupted actin alignment and promoted filament clustering and polymerisation after 48 hours. In contrast, PBM-treated HVPCs displayed a more organised actin architecture, with finer and less aggregated filaments (Figure S7E). This restoration of cytoskeletal organisation may contribute to improved endothelial support and barrier integrity under hypoxic stress. Collectively, these findings highlight distinct, cell-type–specific responses to PBM and suggest that modulation of pericyte cytoskeletal dynamics may represent an additional mechanism through which PBM supports BBB function.

## 4. Discussion

BBB breakdown is increasingly recognised as a central and often early feature of many CNS pathologies, including stroke, traumatic brain injury, and major neurodegenerative disorders, where it contributes to neuroinflammation, metabolic imbalance and progressive neuronal dysfunction. Although it remains debated whether BBB disruption is a primary driver of disease or a secondary amplifier of pathology, targeting BBB dysfunction itself is likely to confer therapeutic benefit across a broad spectrum of brain conditions. In this context, the present study explored a light-driven therapeutic strategy and its potential to restore BBB integrity following hypoxic injury. By modelling the BBB in vitro, we were able to directly assess barrier function and cellular physiology, leading to the identification of PBM-mediated downregulation of vWF as a previously unrecognised mechanism contributing to barrier restoration, alongside modulation of oxidative stress and thrombo-inflammatory signalling pathways.

A central finding of this study is that PBM significantly improves TEER in hypoxia-exposed endothelial monocultures and BBB tri-cultures at both 24 and 48 hours, indicating sustained restoration of barrier function and warranting detailed investigation of the underlying molecular mechanisms. The platelet glycoprotein vWF is released from WPBs into the circulation and is only expressed in endothelial cells and megakaryocytes upon activation. In von Willebrand disease a deficiency of vWF leads to the congenital bleeding disorders, including Bernard-Soulier and platelet-type vWD [21]. Yet, high levels of vWF have been found in plasma from patients with neurological conditions, such as stroke, traumatic brain injuries (TBI), and cerebral malaria [22, 23]. Moreover, various in vivo studies showed the direct regulatory role of vWF on increasing the BBB permeability in mice upon hypoxia/reoxygenation [22] and spontaneous intracerebral haemorrhage [24]. Besides, vWF induced cerebral inflammation, tight junction reorganisation, loss of pericyte coverage and neuronal injury [24]. Both studies demonstrated that genetic deletion of vWF results in a tighter BBB, characterised by reduced permeability, diminished Evans blue extravasation, and lower brain water content. In line with these findings, our data show that PBM significantly downregulates vWF expression in hypoxic endothelial cells, identifying a potential non-invasive strategy to support restoration of BBB integrity under hypoxic stress.

Unbiased profiling of secreted inflammatory mediators using a human cytokine array revealed that PBM significantly reduced the expression of PAI-1 and macrophage MIF in BBB tri-cultures under both normoxic and hypoxic conditions. Similarly to vWF, PAI-1 is involved in the coagulation pathway and inhibits the serine proteases tissue-type plasminogen activator (t-PA) and urokinase plasminogen activator (u-PA), thereby preventing fibrinolysis in the circulation and favouring the formation of blood clots [25]. A recent study showed a reduction in BBB disruption and inflammation after knocking down *SerpinE1* encoding for PAI-1 in mice subjected to ischemic stroke compared to wild-type animals. Furthermore, with PBM acting on two mediators of the thrombotic pathway, red and infrared light might promote the recovery of hypoxia-related neurological conditions by reducing neuro-inflammation via the rescue of the BBB function [26]. Consistent with this, previous studies have shown that modulation of MIF levels attenuates astrocyte activation and cytokine production both in vitro and in vivo, reduces infiltration of peripheral immune cells, and mitigates endothelial cell death and neurological deficits in murine models of perioperative ischaemic stroke [27–30]. Although it is unclear which BBB cell type in the tri-culture primarily downregulated MIF upon PBM, light may not only reduce neuroinflammation but also improve BBB permeability. Liu et al. 2018 showed that tight junction disruption in adult rat brain endothelial cells occurs upon MIF administration and that Evan blue leakage is enhanced after transient middle cerebral occlusion (tMCAo) with MIF. Treating the in vitro and in vivo models with the MIF antagonist ISO-1 restored the ZO-1 disruption, reduced leakage and infarct volume in tMCAo rats, and improved neurological scores [31]. The PBM-mediated downregulation of PAI-1 and MIF could therefore additionally contribute to the improvement in barrier function and TEER after hypoxia. In addition to downregulation, PBM upregulated IL-8 expression in tri-cultures, which has been shown to contribute to the maintenance of endothelial cells, capillary tube organisation, and angiogenesis [32]. Since IL-8 is stored in WPBs and its secretion is controlled by vWF expression, the increase observed in the cytokine dot plot may be initiated by astrocytes and pericytes, highlighting the role of the cell-to-cell contact model [33].

Oxidative stress is a well-established hallmark of hypoxic injury and a key mediator of BBB disruption, contributing to endothelial dysfunction, inflammatory signalling, and reduces cellular viability. Hypoxia did not significantly reduce the cell viability across all cultures compared to normoxia (Figure S1G) and PBM did not alter viability (Figure S1F) or cell cytotoxicity (Figure S1D) compared to non-PBM hypoxic cultures. Yet, upon measuring H_2_O_2_ synthesis after each PBM irradiation of hypoxic cells, this study showed an immediate and pronounced decrease in ROS in astrocytes and pericytes. Thus, PBM’s antioxidant nature could be another pathway supporting the tri-culture function and returning cells into homeostasis [34]. Interestingly, OCR and ECAR data revealed that astrocytes and pericytes were less affected by hypoxia compared to endothelial cells as they show minimal differences between normoxic and hypoxic basal respiration at 24 and 48 hours. Notably, canonical reactivity markers including GFAP, AQP4, and Rgs5, were not significantly upregulated following hypoxic exposure, indicating that under the conditions tested astrocytes and pericytes did not mount a pronounced reactive response. This may reflect intrinsic resilience to transient hypoxia and/or a rapid recovery upon reoxygenation in our model [35, 36]. In contrast, endothelial cells exhibited clear metabolic vulnerability, with hypoxia significantly reducing basal OCR-linked respiration. PBM selectively enhanced maximal respiratory capacity in endothelial cells at 24 hours, suggesting improved mitochondrial reserve and bioenergetic flexibility. Together, these findings support the interpretation that PBM-mediated restoration of barrier function is primarily driven by endothelial rescue rather than glial reprogramming. By directly improving endothelial metabolic competence and attenuating hypoxia-induced dysfunction, red and near-infrared light therapy emerges as a strategy specifically targeting vascular health, with broader implications for conditions characterised by endothelial instability and impaired neurovascular coupling. Another possible pathway involved downregulation of HIF-1α, as detected by ICC. This link has already been discussed by Engelhardt et al. (2013), who reported that HIF-1α stabilisation in hypoxia-or ischemia-induced injuries is closely associated with BBB dysfunction, and that inhibition improves the barrier integrity of brain endothelial cells [14, 37]. Thus, modulating HIF-1α, ROS levels, and mitochondrial function could support the maintenance of tight junctions and barrier integrity following hypoxia.

Although the transwell system does not incorporate physiological shear stress or tubular geometry characteristic of 3D microvascular models, it provides a highly controlled and reproducible platform for isolating cell-specific molecular responses and quantifying barrier integrity with precision. For the mechanistic questions addressed here - namely the direct effects of hypoxia and PBM on endothelial-led barrier recovery and multicellular signalling - this level of experimental control was essential. The use of immortalised human cell lines further ensured consistency across replicates, enabling robust statistical comparisons and clear attribution of molecular changes to defined cell populations.

We recognise that incorporation of flow-based systems, primary or iPSC-derived cells, and immune components such as microglia would add additional physiological complexity and allow deeper interrogation of inflammatory crosstalk and long-term remodelling. These represent logical next steps building directly on the mechanistic framework established here. Similarly, while hypoxia is a broadly applicable insult, it models a convergent pathway common to stroke, traumatic brain injury, and chronic neurodegenerative disease, making the findings broadly relevant across CNS pathologies.

Importantly, this study establishes a clear molecular link between PBM and restoration of BBB function under hypoxic stress, identifying endothelial vWF regulation and coordinated oxidative and thrombo-inflammatory modulation as actionable targets. In a therapeutic landscape where options to directly stabilise the BBB remain limited and often associated with systemic adverse effects, PBM offers a non-invasive, mechanistically grounded strategy for vascular protection [38].

## 5. Conclusion

In conclusion, this study establishes a mechanistically anchored in vitro framework to dissect how hypoxia disrupts BBB integrity and how targeted PBM modulates this response as outlined in Figure 6. Within a multicellular human BBB model, endothelial cells emerged as the primary drivers of barrier dysfunction under hypoxic stress, exhibiting marked alterations in TEER, oxidative status and secretory phenotype. Importantly, PBM reproducibly improved barrier integrity and attenuated endothelial activation, with complementary genetic perturbation experiments identifying endothelial vWF as a functionally relevant mediator of barrier recovery.

**Figure 6:**
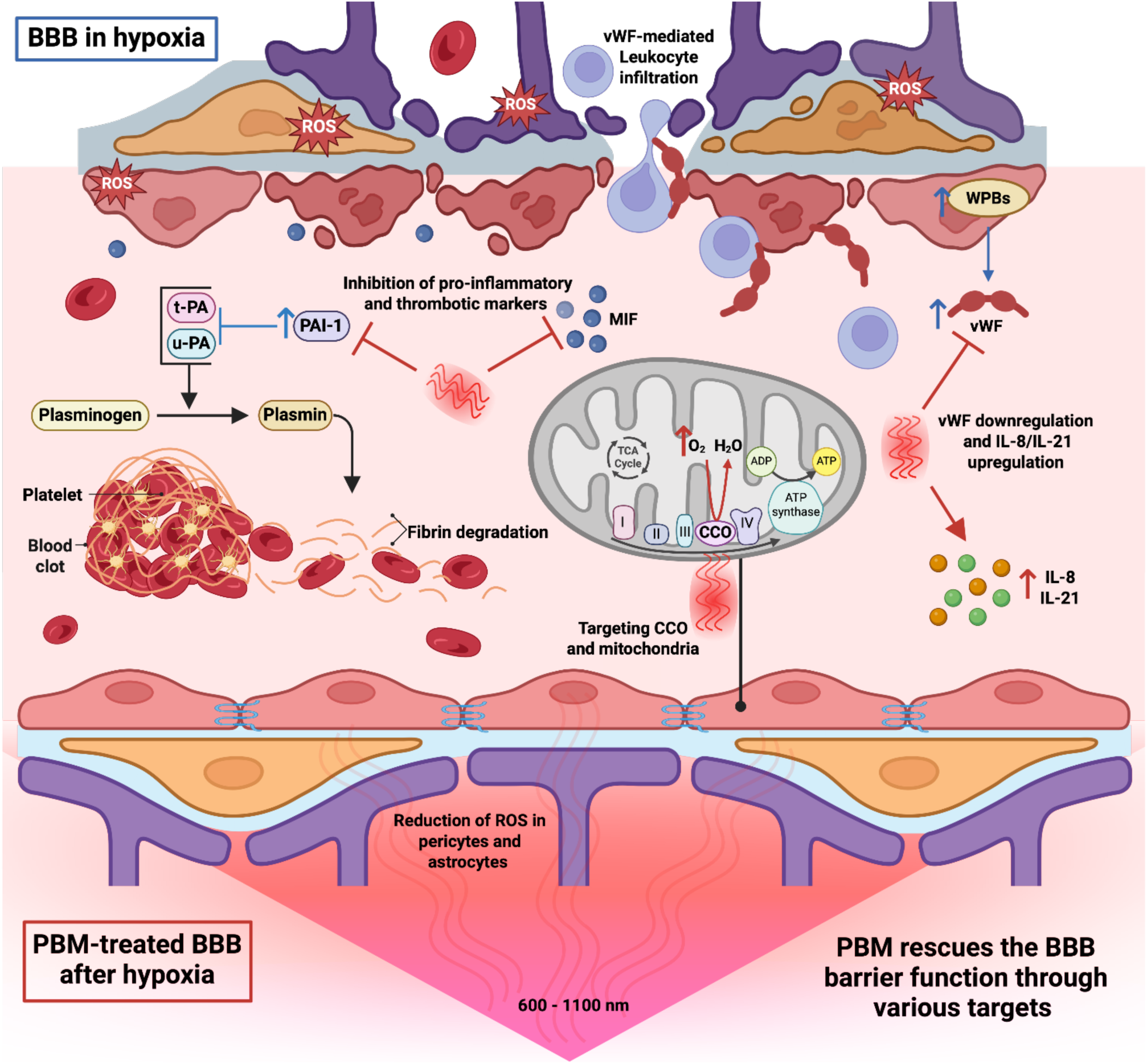
Schematic drawing of molecular targets of PBM of the BBB after hypoxia that led to the rescue of barrier function. Created with Biorender.

By combining barrier physiology, mitochondrial bioenergetics and targeted molecular manipulation, this work moves beyond descriptive observation and identifies a discrete endothelial node linking hypoxic activation to barrier failure. While conducted in an in vitro system, the model allows controlled interrogation of cell-type–specific mechanisms that are difficult to resolve in vivo. The data therefore provide both mechanistic insight and a tractable platform for future validation in primary and translational settings.

Collectively, these findings position PBM not simply as a broadly protective stimulus, but as a modulator of defined endothelial pathways that regulate barrier integrity. This work strengthens the conceptual framework linking hypoxia, endothelial activation and BBB dysfunction, and identifies actionable molecular targets for therapeutic exploration in acute brain injury and in chronic disorders where vascular compromise contributes to progressive neurodegeneration.

## Authors’ translational perspective

BBB dysfunction is increasingly recognised as a central contributor to neurological injury and disease progression, yet therapeutic strategies that directly target barrier repair remain limited. Our findings suggest that PBM can restore endothelial barrier integrity after hypoxic stress by modulating defined thrombo-inflammatory and metabolic pathways, including HIF-1α signalling and vWF expression. Importantly, the data indicate that endothelial cells represent the primary responders to hypoxia within the neurovascular unit, and that targeted modulation of endothelial activation may be sufficient to stabilise barrier function.

While our work is based on an in vitro human BBB model, the mechanistic insights provide a rational framework for further validation in more complex systems, including organotypic models and in vivo settings. The identification of vWF as a functionally relevant mediator of barrier rescue highlights a pathway that could be explored pharmacologically in hypoxia-associated conditions such as stroke, traumatic brain injury, and chronic cerebrovascular insufficiency.

PBM is non-invasive and has an established safety profile in other clinical contexts. Our data therefore support the concept that carefully timed modulation of endothelial signalling could complement existing neuroprotective strategies. Future work should determine optimal dosing paradigms, depth of light penetration in vivo, and durability of barrier rescue to evaluate whether this approach can meaningfully translate into clinical benefit.

## Additional information

### Data and materials availability

All data needed to evaluate the conclusions in the paper are present in the paper and/or the Supplementary Materials. This study did not generate new unique reagents.

### Author contributions

M.M.S conceived the study and devised the experimental strategy with L.S.

M.M.S and L.S. supervised the research. M.D. performed all the experiments and analysed the data with support from L.S. D.E.B and N.S provided critical knowledge to this work. M.D. assembled figures. M.D., L.S. and M.M.S wrote the manuscript. All authors edited, read, and approved the manuscript.

### Funding and acknowledgements

The authors would like to thank Joe DiDuro, Sam Larson and Roslyn Bill for thoughtful discussions on the study. M.M.S., L.S. and M.D. are supported by a Medical Research Council Career Development Award (MR/W027119/1) and by the British Heart Foundation and the UK Dementia Research Institute (award number UK DRI-8203) through UK DRI Ltd, principally funded by the Medical Research Council. M.M.S acknowledges support from the BHF Centre of Research Excellence, University of Oxford (grant code: RE/24/130024). L.S. acknowledges the support by the Royal Society Newton International Fellowship (NIF\R1\242594).

## Abbreviations

2-DG: 2-Deoxy-Glucose
αSMA: Alpha smooth muscle actin
AM: Astrocyte medium
ATP: Adenosine triphosphate
AQP4: Aquaporin-4
BBB: Blood-brain barrier
BS: Blocking solution
CCO: Cytochrome C oxidase
CNS: Central nervous system
dH_2_O: Distilled water
ECAR: Extracellular acidification rate
EM: Endothelial cell medium
FCCP: Carbonyl cyanide-4 (trifluoromethoxy) phenylhydrazone
FDA: Food and drug administration
eNO: Endothelial nitric oxide
GFAP: Glial fibrillary acidic protein
GM-CSF: Granulocyte-macrophage colony-stimulating factor
GROα: Growth-regulated oncogene-α
HAs: Human astrocytes
HK: Housekeeping
H_2_O_2_: Hydrogen peroxide
HBMECs: Human brain microvascular endothelial cells
HVPCs: Human brain vascular pericytes
HIF-1α: Hypoxia-inducible factor-1 alpha
ICC: Immunocytochemistry
IL-8: Interleukin-8
IL-21: Interleukin-21
LLLT: Low level laser therapy
MIF: Migration inhibitory factor
O_2_: Oxygen
OCR: Oxygen consumption rate
PBM: Photobiomodulation
PDGFRβ: Platelet-derived growth factor receptor-β
PFA: Paraformaldehyde
PLL: Poly-L-lysine
PM: Pericyte medium-PLUS
qPCR: quantitative polymerase chain reaction
R/A: Rotenone/Antimycin A
Rgs5: Regulator of G-protein signalling 5
RLU: Relative average luminescence
RPL13A: Ribosomal protein 13A
ROS: Reactive oxygen species
SEM: Standard error of the mean
Serpin E1/PAI-1: Plasminogen activator inhibitor-1
siRNA: Small interfering RNA
TBI: Traumatic brain injury
TEER: Transendothelial electrical resistance
TGF-β: Transforming growth factor beta
t-PA: Tissue-type plasminogen activator
u-PA: Urokinase-type plasminogen activator
v-vWF: von Willebrand factor
WPBs: Weibel-Palade bodies
ZO-1: Zonula occludens-1

## Supplementary Tables

**Table S1.** The relative expression of positive and not significant cytokines, chemokines, and acute phase proteins for all four conditions.

| Cytokine | Normoxia | Normoxia + PBM | Hypoxia | Hypoxia+PBM |
| --- | --- | --- | --- | --- |
| CCL2/MCP-1 | 1 | 1.1404 | 0.84415 | 0.99055 |
| CXCL12/SDF-1 | 1 | 1.1404 | 1.0444 | 0.7455 |
| G-CSF | 1 | 1.04675 | 0.5955 | 0.7737 |
| ICAM-1/CD54 | 1 | 1.1677 | 0.7578 | 0.90745 |
| IL-2 | 1 | 1.0617 | 0.49415 | 0.46825 |
| IL-6 | 1 | 1.23435 | 0.81775 | 0.9144 |
| IL-13 | 1 | 0.8251 | 0.5311 | 0.32685 |
| IL-16 | 1 | 0.8036 | 0.5099 | 0.26735 |
| IL-17A | 1 | 1.0614 | 0.53025 | 0.26835 |
| IL-18/IL-1F4 | 1 | 1.03755 | 0.78855 | 0.7347 |
| IL-32α | 1 | 0.96455 | 0.4543 | 0.21215 |

**Table S2.** The cell counts per biological replicate used for the transwell model of WT and siRNA treated endothelial cells. The mean of the cell count between WT and transfected endothelial cells was compared with an unpaired two-tailed t-test with N=4 (biological replicates).

| Endothelial cell count | WT (x10<sup>6</sup> cells/mL) | siRNA (x10<sup>6</sup> cells/mL) | Significance |
| --- | --- | --- | --- |
| N=1 | 6.33 | 4.57 | - |
| N=2 | 3.96 | 3.77 | - |
| N=3 | 5.60 | 5.06 | - |
| N=4 | 5.63 | 4.70 | - |
| Mean +/- SEM | 5.380 +/- 0.5024 | 4.525 +/- 0.2722 | ns (p=0.0851) |

## Supplementary Figures

**Figure S1.**
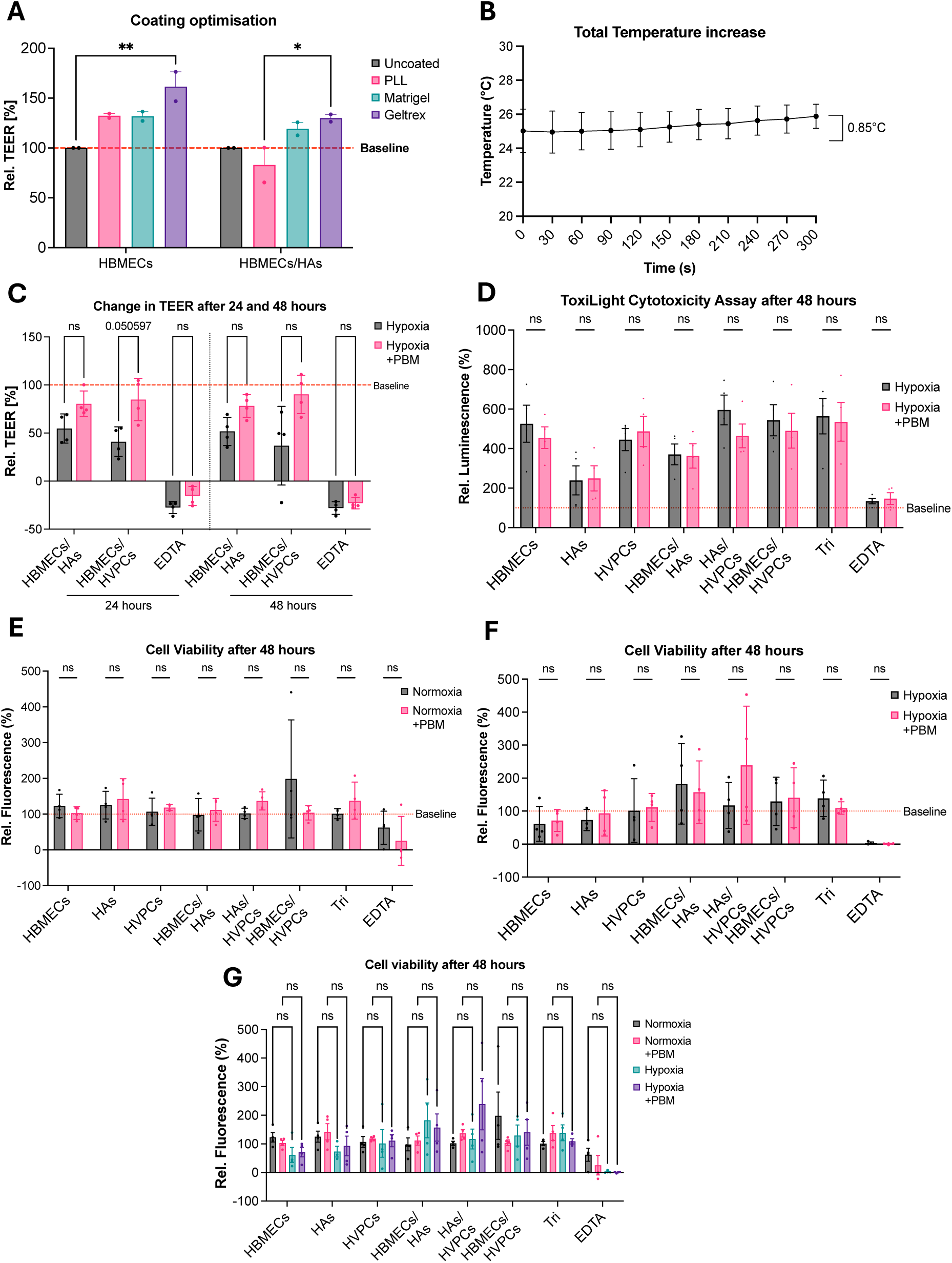
Validation of the transwell BBB model and assessment of PBM-related controls. (A) Relative change in TEER (%) on day 8 in endothelial monocultures and endothelial–astrocyte co-cultures under different apical coating conditions. Means were analysed by two-way ANOVA with Tukey’s multiple comparisons test (*p<0.05, **p<0.01; n=2 technical replicates from one biological experiment). (B) Temperature changes recorded in the transwell supernatant during a single PBM exposure, confirming the absence of thermal effects. (C) Relative change in TEER (%) in co-cultures that did not show significant barrier alterations 24 and 48 h after hypoxia. (D) Relative luminescence signal (%) assessing cytotoxicity across all transwell conditions (±PBM) 48 h after hypoxia. (E–F) Relative fluorescence signal (%) assessing cell viability in normoxia (E) and 48 h after hypoxia (F) across all transwell conditions (±PBM). (G) Relative fluorescence signal (%) comparing normoxia and hypoxia at 48 h. For panels C–G, means were analysed using multiple unpaired t-tests with Holm–Šídák correction (N=4 biological replicates). Data are presented as mean ± SEM.

**Figure S2.**
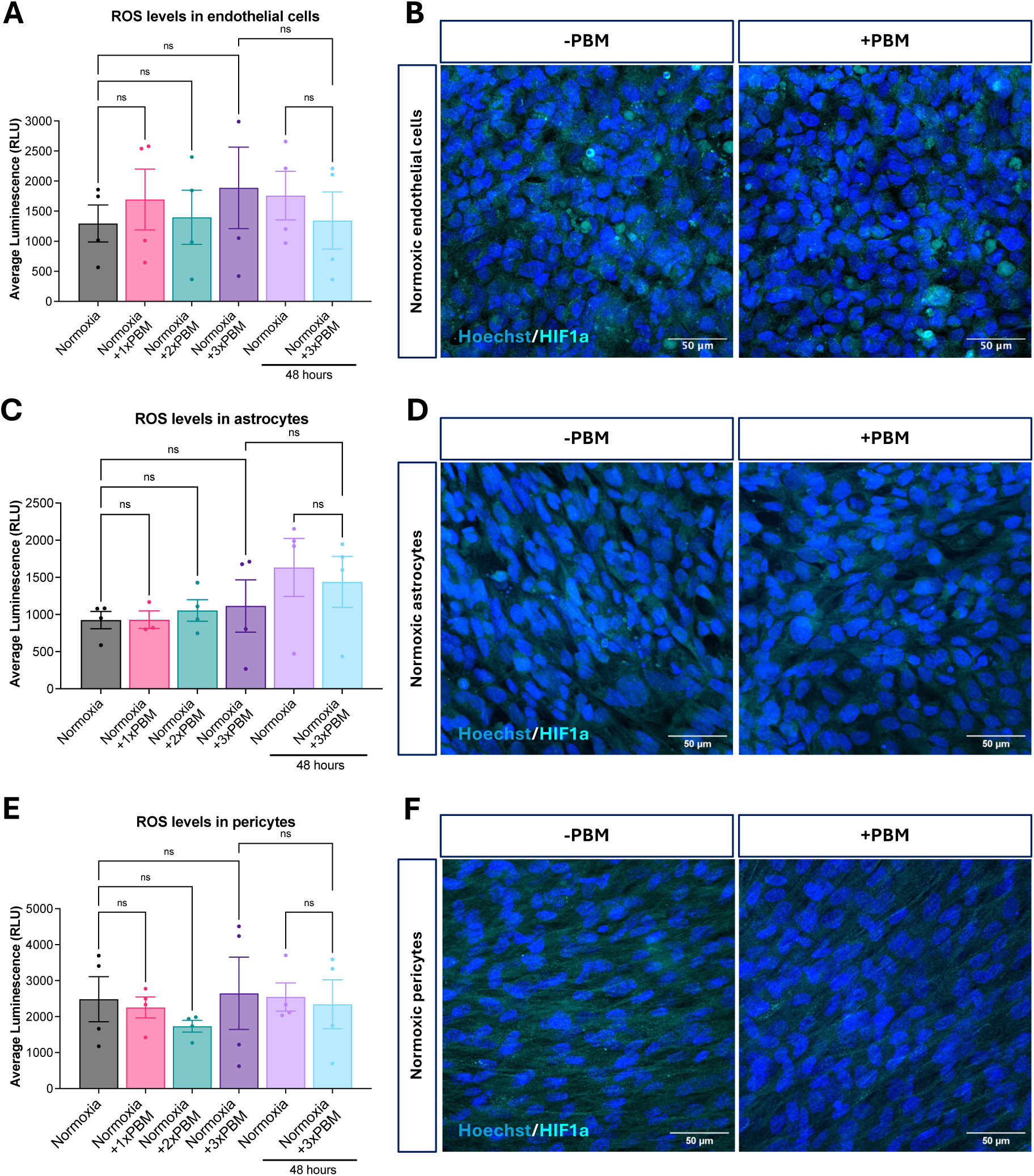
PBM does not significantly alter ROS or HIF-1α under normoxic conditions. (A,C,E) Average luminescence signal quantifying ROS levels in normoxic endothelial cells (A), astrocytes (C), and pericytes (E), comparing untreated and PBM-treated groups. (B,D,F) Representative ICC images of HIF-1α (cyan) with Hoechst nuclear counterstain (blue) in normoxic endothelial cells (B), astrocytes (D), and pericytes (F) under non-PBM (–PBM) and PBM-treated (+PBM) conditions. Quantitative comparisons were performed using one-way ANOVA with Šidák’s multiple comparisons test (N=4 biological replicates). Data are presented as mean ± SEM.

**Figure S3.**
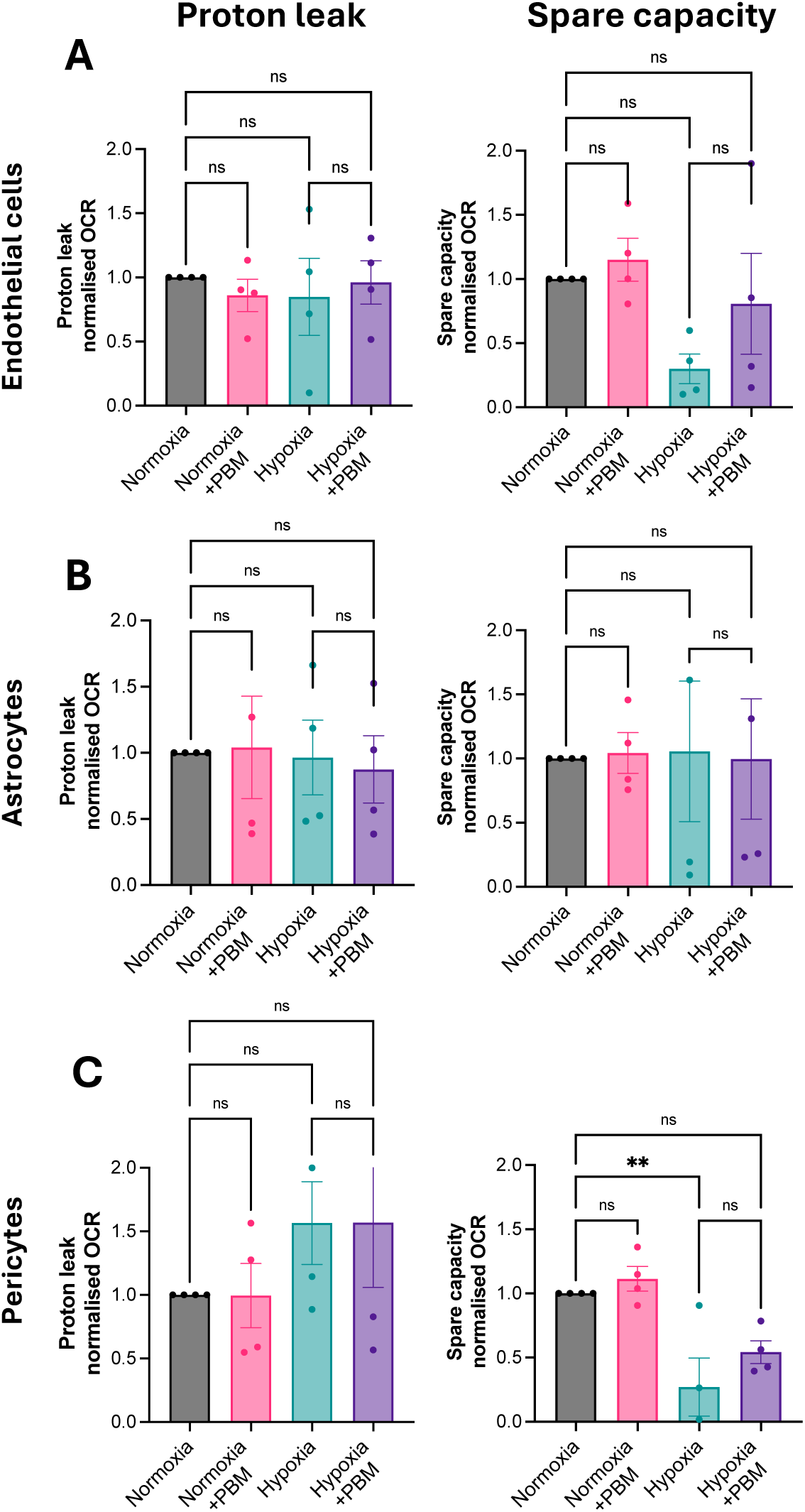
PBM effects on mitochondrial proton leak and spare respiratory capacity following hypoxia. (A-C) Quantification of proton leak and spare respiratory capacity 24 hours after treatment in endothelial cells (A), astrocytes (B), and pericytes (C). Proton leak was calculated from OCR values following oligomycin injection and normalised to total OCR. Spare respiratory capacity was determined as the difference between maximal respiration (post-FCCP) and basal respiration, normalised to OCR. Statistical comparisons were performed using one-way ANOVA with Šidák’s multiple comparisons test (N=4 biological replicates; **p<0.01). Data are presented as mean ± SEM.

**Figure S4.**
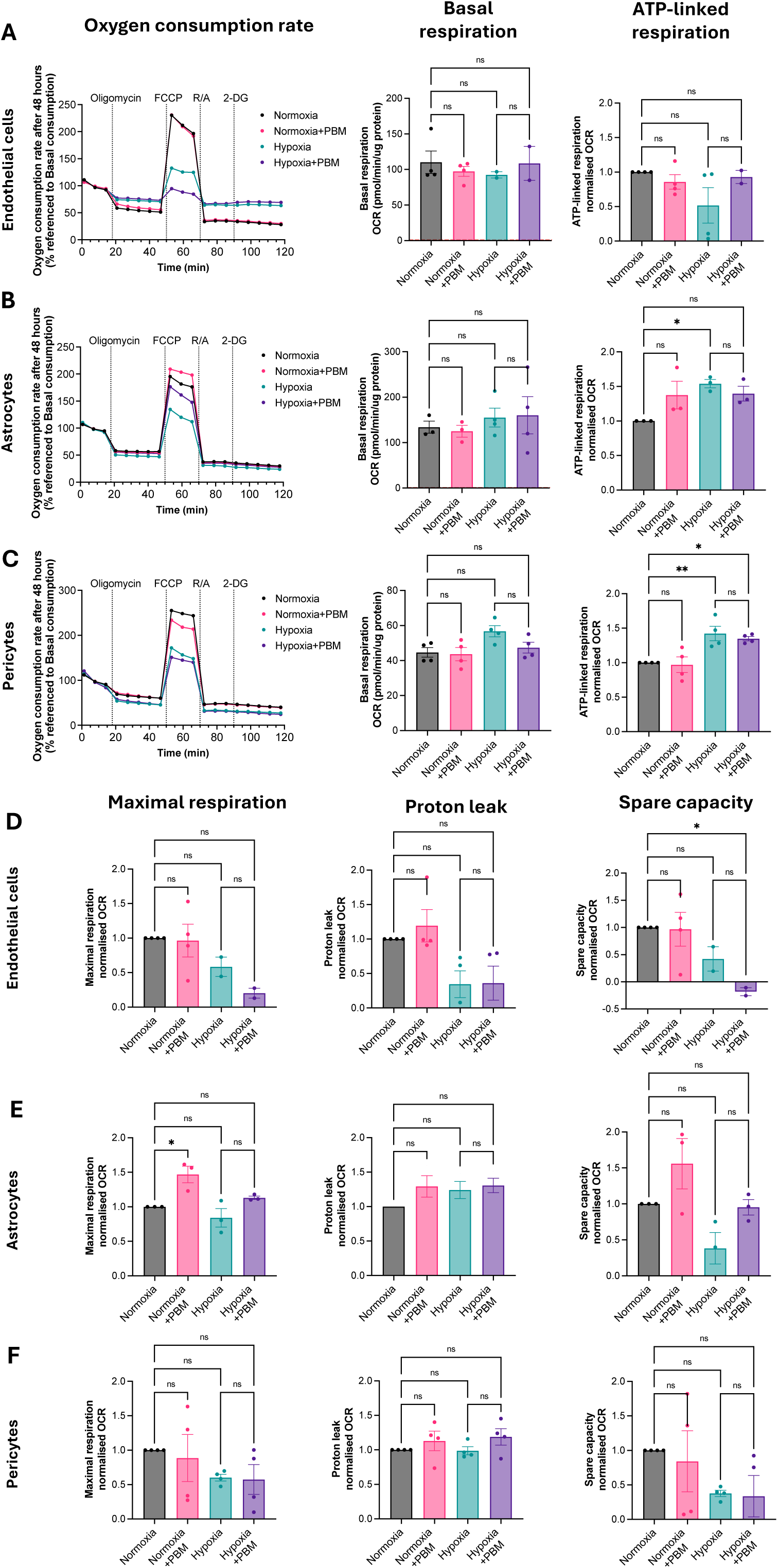
PBM influences on mitochondrial respiratory parameters 48 hours after hypoxic injury. (A–F) Oxygen consumption rate (OCR) measurements in endothelial cells (A,D), astrocytes (B,E), and pericytes (C,F) 48 hours post-treatment. OCR traces are plotted over time and normalised to protein content and baseline respiration. Basal respiration (pmol/min/µg protein) was calculated prior to oligomycin injection. ATP-linked respiration was determined as the difference between basal respiration and OCR following oligomycin, normalised to total OCR. Maximal respiration was calculated from OCR after FCCP injection and normalised accordingly. Proton leak was derived from OCR after oligomycin injection, and spare respiratory capacity was defined as the difference between maximal and basal respiration, both normalised to OCR. Statistical comparisons were performed using one-way ANOVA with Šidák’s multiple comparisons test (N=4 biological replicates; *p<0.05, **p<0.01). Data are presented as mean ± SEM.

**Figure S5.**
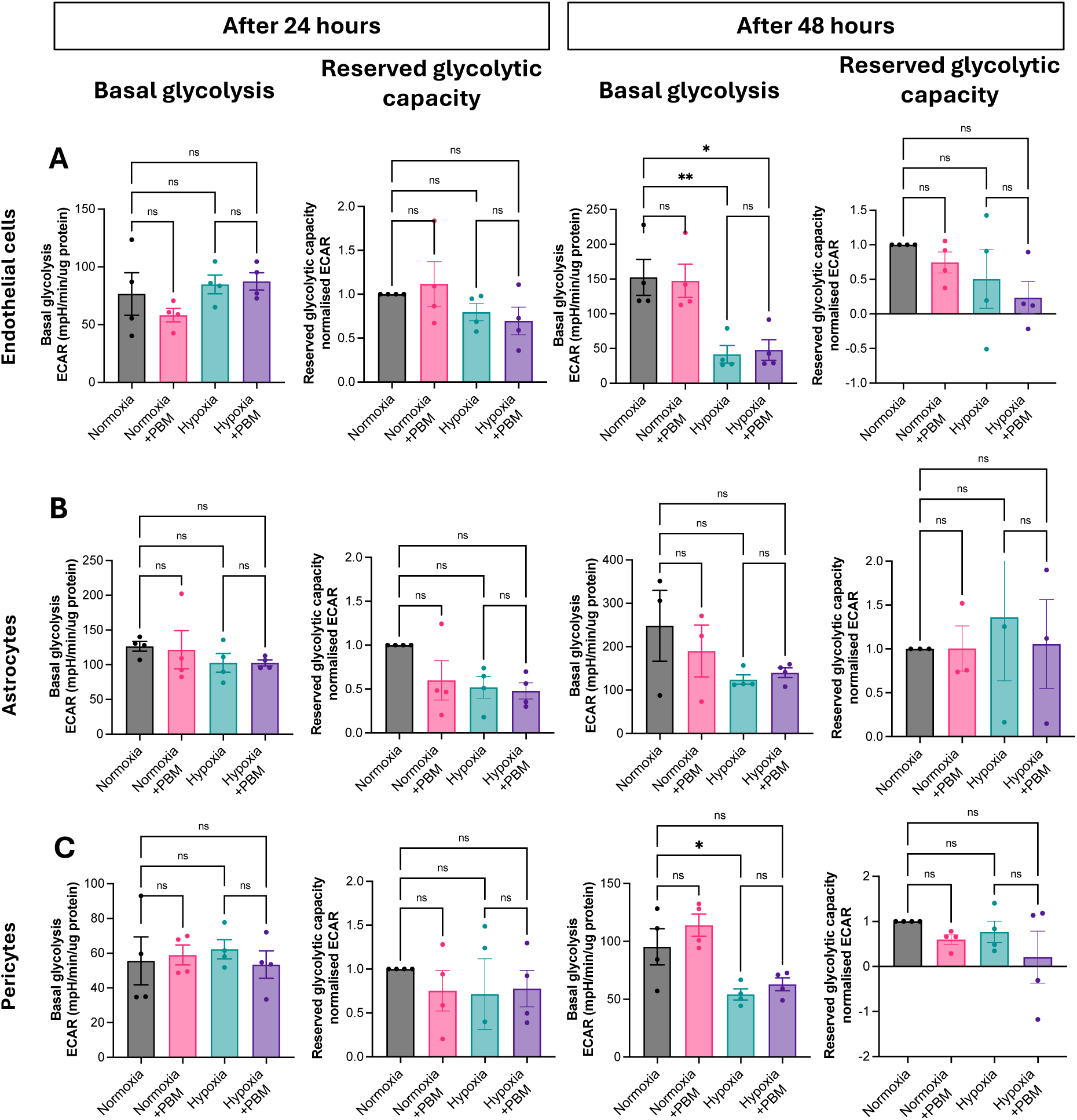
PBM effects on glycolytic parameters following hypoxic exposure after 24 and 48h. (A–C) ECAR measurements in endothelial cells (A), astrocytes (B), and pericytes (C) at 24 and 48 hours across all experimental conditions. Basal glycolysis (mpH/min/µg protein) was calculated prior to oligomycin injection. Glycolytic reserve capacity was defined as the difference between ECAR following oligomycin and basal glycolysis, normalised to total ECAR. Statistical comparisons were performed using one-way ANOVA with Šidák’s multiple comparisons test (N=4 biological replicates; *p<0.05, **p<0.01). Data are presented as mean ± SEM.

**Figure S6.**
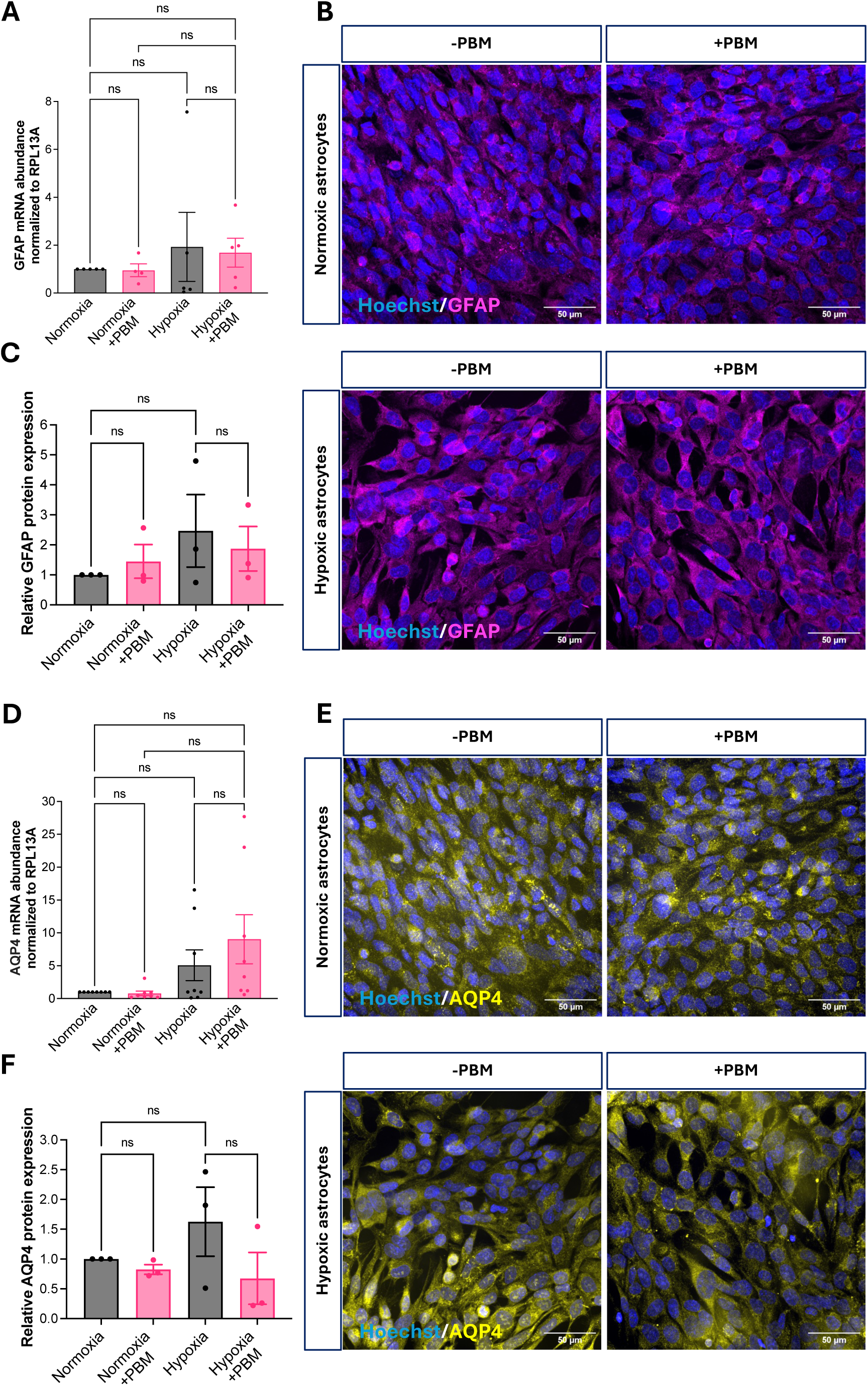
Astrocytic GFAP and AQP4 expression following hypoxia and PBM treatment. (A) GFAP mRNA expression in astrocytes at 48 hours across all experimental conditions, normalised to RPL13A. (B) Representative ICC images showing GFAP (magenta) and nuclei (Hoechst, blue) in normoxic and hypoxic astrocytes, with and without PBM treatment. (C) Quantification of GFAP protein expression relative to control, calculated as mean fluorescence intensity normalised to Hoechst area. (D) AQP4 mRNA expression in astrocytes at 48 hours, normalised to RPL13A. (E) Representative ICC images showing AQP4 (yellow) and nuclei (blue) under normoxic and hypoxic conditions ± PBM. (F) Quantitative analysis of AQP4 protein expression relative to control, measured as mean fluorescence intensity normalised to Hoechst area. Statistical comparisons were performed using one-way ANOVA with Šidák’s multiple comparisons test (N=3–4 biological replicates). Data are presented as mean ± SEM.

**Figure S7.**
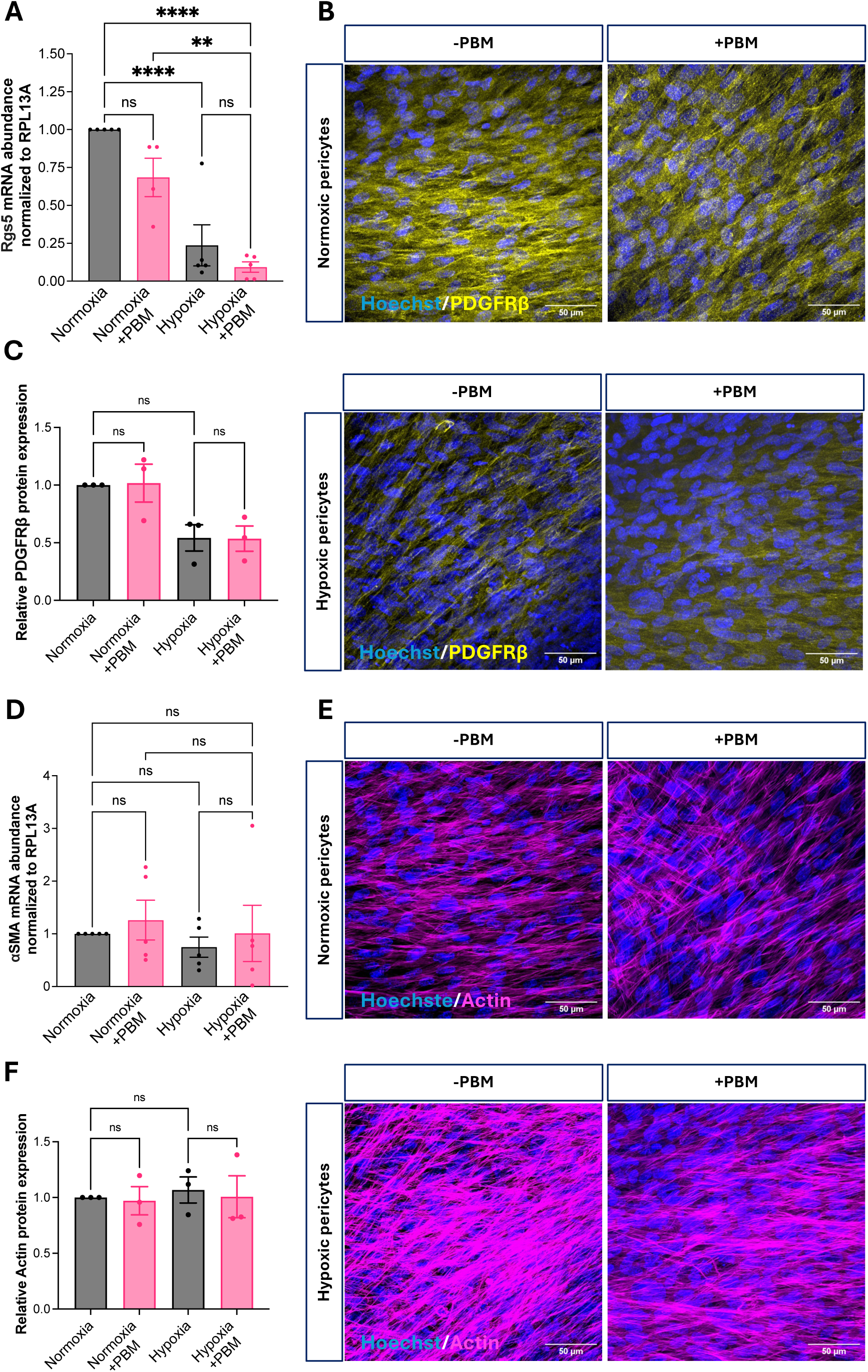
Pericyte-specific responses: PDGFRβ, Rgs5, αSMA and actin organisation following hypoxia and PBM. (A) Rgs5 mRNA expression in human vascular pericytes (HVPCs) at 48 hours across all experimental conditions, normalised to RPL13A. (B) Representative ICC images showing PDGFRβ (yellow) and nuclei (Hoechst, blue) in normoxic and hypoxic HVPCs, with and without PBM treatment. (C) Quantification of PDGFRβ protein expression relative to control, calculated as mean fluorescence intensity normalised to Hoechst area. (D) αSMA mRNA expression in HVPCs at 48 hours, normalised to RPL13A. (E) Representative ICC images showing filamentous actin (magenta) and nuclei (blue) under normoxic and hypoxic conditions ± PBM. (F) Quantitative analysis of actin protein expression relative to control, measured as mean fluorescence intensity normalised to Hoechst area. Statistical comparisons were performed using one-way ANOVA with Šidák’s multiple comparisons test (N=3–4 biological replicates). Data are presented as mean ± SEM.

